# Low cell number proteomic analysis using in-cell protease digests reveals a robust signature for cell cycle state classification

**DOI:** 10.1101/2020.07.03.186023

**Authors:** Kelly Van, Aymen al-Rawi, David Lewis, Georg Kustatscher, Tony Ly

**Affiliations:** Wellcome Centre for Cell Biology, School of Biological Sciences, University of Edinburgh, Michael Swann Building, Max Born Crescent, Edinburgh EH9 3BF, UK; Institute of Quantitative Biology, Biochemistry and Biotechnology, University of Edinburgh, Michael Swann Building, Max Born Crescent, Edinburgh EH9 3BF, UK; Centre for Gene Regulation and Expression, School of Life Sciences, University of Dundee, Dow Street, Dundee, DD1 5EH

**Author notes:** Correspondence should be addressed to Tony Ly.

## Abstract

Comprehensive proteome analysis of rare cell phenotypes remains a significant challenge. We report a method for low cell number mass spectrometry (MS)-based proteomics using protease digestion of mildly formaldehyde-fixed cells *in cellulo*, which we call the ‘in-cell digest’. We combined this with AMPL (Averaged MS1 Precursor Library Matching) to quantitatively characterise proteomes from low cell numbers of human lymphoblasts. 4,500 proteins were detected from 2,000 cells and 2,500 proteins were quantitated from 200 lymphoblasts. The ease of sample processing and high sensitivity makes this method exceptionally suited for the proteomic analysis of rare cell states, including immune cell subsets and cell cycle subphases.

To demonstrate the method, we characterised the proteome changes across 16 cell cycle states isolated from an asynchronous TK6 human lymphoblast culture, avoiding synchronization. States included late mitotic cells present at extremely low frequency. We identified 119 pseudoperiodic proteins (PsPs) that vary across the cell cycle. Clustering of the PsPs showed abundance patterns consistent with ‘waves’ of protein degradation in late S, at the G2&M border, mid-mitosis and at mitotic exit. These clusters were distinguished by significant differences in predicted nuclear localization and interaction with the APC/C. The dataset also identifies putative APC/C substrates in mitosis and the temporal order in which they are targeted for degradation.

We demonstrate that a protein signature made of these 119 high confidence cell cycle regulated proteins can be used to perform unbiased classification of proteomes into cell cycle states. We applied this signature to 296 proteomes that encompass a range of quantitation methods, cell types, and experimental conditions. The analysis confidently assigns a cell cycle state for 49 proteomes, including correct classification for proteomes from synchronized cells. We anticipate this robust cell cycle protein signature will be crucial for classifying cell states in single cell proteomes.

## Introduction

The proteome is a functional readout of cellular phenotype. Cellular phenotype includes dynamic and persistent features that reflect cell state and cell type, respectively. Rare cell phenotypes play key physiological roles. Quiescent stem cells, while often rare relative to differentiated cell types in a tissue, are essential for tissue homeostasis. Similarly, mitosis is a dynamic cell state that is critical for the accurate propagation of genetic material. Mitotic states are generally short-lived and thus rare in an asynchronous population. Proteomic analysis of these critically important cell phenotypes is a major challenge because typical proteomic workflows require >10^5^ cells as input.

We previously developed an approach called ‘PRIMMUS’ or ‘*P*roteomics of *I*ntracellular I*mmu*nostained *Subsets*” to analyse abundant and rare cell cycle states [1]. Formaldehyde-fixed cells are fractionated into specific cell states by staining cells for intracellular markers and separating them using Fluorescence-Activated Cell Sorting (FACS). Cells grown in asynchronous culture are immediately fixed, thereby minimizing perturbation to physiological processes. This step is critical, as small molecule-based synchronsation can lead to effects on the proteome that are associated with stress responses arising from arrest rather than cell cycle regulation *per se* [2]. PRIMMUS enabled analysis of interphase and mitotic subpopulations, but this approach was limited to relatively abundant subpopulations for which >10^5^ cells can be collected by FACS within a reasonable time [3].

Low input proteome analysis requires specialized methods for handling low cell number of cells [4,5]. Major improvements have been made by adapting methods used for bulk samples to low cell number samples [6-8]. Recent advances in small volume sample handling to nanoliter volumes have also enabled analysis of <10 cultured human cells, with overall number of proteins detected scaling with cell number [4,9,10]. For example, ∼3,000 proteins were identified from 10 HeLa cells using ‘nanodroplet processing in one pot for trace samples’ [11]. In general, these methods require specialized equipment, ranging from automated robotic sample handling to custom microfabricated chips, which are challenging to satisfy in most labs currently.

Cells fixed with formaldehyde introduces additional challenges for bottom-up MS-based proteomics. Formaldehyde crosslinks proteins by forming methylene bridges primarily between lysine residues. Peptide/protein crosslinks are broken with heating to >65 °C. As an example, formalin fixed tissue processing protocols include heating for 1 hour at 95 °C. However, the fixative concentration and treatment duration for formalin fixed tissues is much higher (4% formaldehyde for up to several hours). Studies on synthetic peptides demonstrated that protein amino acid residues can be irreversibly modified by formaldehyde, producing chemical modifications and corresponding mass shifts that are not detected in conventional database searches [12,13]. In contrast, formaldehyde fixation for immune cell immunostaining and flow cytometry in clinical and academic research settings is frequently much lower (0.1 – 3%) and carried out under controlled conditions with limited treatment duration (10 – 30 min).

Here, we report a methodological advance that eliminates several steps previously required for processing fixed cells for proteomics. We demonstrate that fixed cells in suspension can be directly digested by trypsin without heat-induced crosslink reversal for quantitative proteomics. We call this streamlined approach the *in-cell digest*. The in-cell digest provides major improvements in sensitivity and convenience in performing proteomic analysis on low numbers of fixed cells. To overcome the duty cycle limitations of the Orbitrap Elite instrument, we developed an acquisition method called AMPL. We applied in-cell digest and AMPL with PRIMMUS to analyse the proteomic variation during an unperturbed cell cycle in human lymphoblasts with unparalleled temporal resolution to produce unbiased proteomic definitions of cell cycle state.

## Experimental Procedures

### Experimental Design and Statistical Rationale

Four biological replicates of 16 cell cycle populations were collected by FACS, with 2 technical replicates of the 64 samples being acquired by LCMS (AMP acquisition strategy) resulting in 128 LCMS analyses, providing 8 pseudotimecoures for periodicity analysis. Three libraries were generated from 12 HPRP fractions of unsorted cells, interphase cells, and mitotic cells. Each library fraction was analysed twice (or thrice for the mitotic library) resulting in a library of 85 LCMS analyses. Libraries were used to increase proteome coverage through MS1 feature matching.

Supporting experiments include the analysis of 12 HPRP fractions of formaldehyde fixed, fixed and reversed, and non-fixed control without replicates for a qualitative comparison of peptide modifications. Twelve cell titration samples were also collected in duplicate up to 2,000 cells by FACS, including a zero-cell control, to assess LC-MS sensitivity of the improved processing and AMP acquisition methods. The 24 cell titration samples were analysed by AMP LCMS along with a 12 HPRP fraction library and an unfractionated library of 2000 sorted cells, which were analysed by DDA LC-MS. To assess the impact of peptide filtering on MS1 feature matching FDR, unmodified, dimethylated, and isopropylated peptides were analysed by AMPL and DDA, along with a library of 12 HPRP fractions.

### Cell culture

TK6 human lymphoblasts [14] were obtained from the Earnshaw laboratory (University of Edinburgh). Cells were cultured at 37 °C in the presence of 5% CO_2_ as a suspension in RPMI-1640 + GlutaMAX (Thermo Scientific) supplemented with 10% v/v fetal bovine serum (FBS, Thermo Scientific). Cell cultures were maintained at densities no higher than 2 x 10^6^ cells per ml. MCF10A cells (ATCC) were cultured in phenol red-free F12/DMEM media (Thermo Scientific) supplemented with 5% horse serum, 10 µg/ml insulin (Sigma), 100 ng/ml cholera toxin (Sigma), 20 ng/ml EGF (Sigma), 0.5 µg/ml hydrocortisone (Sigma), 100 units/ml penicillin and 100 µg/ml streptomycin (Thermo Scientific) at 37 °C in the presence of 5% CO_2_. Cells were maintained at less than 100% confluency and were discarded when passage number exceed 20 passages. U2OS cells (ATCC) were cultured in DMEM media high glucose + GlutaMAX (Thermo Scientific) supplemented with 10% v/v FBS (Thermo Scientific). Cells were checked for mycoplasma at the point of cryo-storage using a luminescence-based assay (Lonza).

### Cell fixation and immunostaining

Cells were washed with Dulbecco’s phosphate-buffered saline (DPBS, Lonza) and resuspended in freshly prepared 1% formaldehyde solution (w/v) from a 16% stock (w/v, Thermo Scientific) in DPBS, fixed for 10 min at room temperature with gentle rotation, pelleted, washed with DPBS and permeabilized with cold 90% methanol. Cells were stored at −20°C prior to staining.

Cells stored in methanol were washed with DPBS and resuspended in blocking buffer, which is composed of 5% bovine serum albumin (BSA) in 0.1 M Tris-buffered saline, pH 7.4 (TBS). Cells were blocked for 10 min at room temperature, pelleted, and resuspended in primary antibody solution. The rat anti-H3S28ph HTA28 (abcam ab10543), mouse anti-cyclin A2 (Cell Signaling Technologies 4656S), and rabbit anti-cyclin B1 (12231S) were used for staining as 1:200 dilutions in blocking buffer. Cells were stained with primary antibody overnight at 4°C. Stained cells were then washed twice with wash buffer (DPBS + 0.5% BSA) and stained with dye-conjugated secondary antibodies. The donkey anti-rat IgG H&L AlexaFluor 568 preadsorbed (abcam ab175475), Donkey anti Mouse IgG (H+L) Highly Cross Adsorbed Secondary Antibody, Alexa Fluor 488 (Thermo A21202), and Goat Anti-Rabbit IgG H&L (Alexa Fluor® 647) preadsorbed (abcam ab150083) were used as 1:200 dilutions in blocking buffer. Cells were stained stained for 1 hour at room temperature, washed twice with DPBS, pelleted, and stained in DAPI solution (Sigma, 20 µg/ml in DPBS + 0.1% BSA) for at least 1 hour prior to FACS.

### FACS and gating strategy

Cells were collected using a BD FACSAria Fusion Cell Sorter equipped with 355nm UV, 405nm Violet, 488nm Blue, 561nm YG and 640nm Red lasers, and controlled by BD FACS Diva V8.0.1 software. Cells were first gated into ‘narrow’ (P1 – P8) and ‘wide’ (P9 – P16) populations based on DAPI fluorescence signal width. The narrow population contains single cells either in interphase, or in mitosis up to late anaphase. These single cells were then separated based on cyclin B into 8 different stages of interphase.

Population P1 has low to no cyclin B protein and 2N DNA content, consistent with low to no E2F activity and a G0/early G1 cell state. Cyclin B rises monotonically from P2 to P6 and then rises more steeply from P6 to P8. Like cyclin B, cyclin A also increases during interphase, but at a faster rate from P1 to P6 as compared to P6 to P8. P9 to P13 are positive for histone H3 phosphorylation at Ser28 (pH3+). Highest levels of pH3+ are present in prometaphase and metaphase. Rising and declining H3 phosphorylation in early and late mitosis, respectively, result in low to medium levels of pH3+. Cyclin A and cyclin B levels are used to further discriminate mitotic subphases, as they are degraded during prometaphase and the metaphase-to-anaphase transition, respectively.

Finally, late mitotic subphases are enriched in the wide population, but so too are doublets. We reasoned that most doublets will have cyclin B signal, as single cells with the exception of P1 are cyclin B positive. Thus, we can further enrich late mitotic stages by selecting wide, 4N, cyclin B negative cells (P14-P16). P14-P16 are then discriminated further by pH3+ levels, which decrease during mitotic exit. We note that P16 may contain doublets of G0/early G1 cells (P1), but P14 and P15 should not as P14 and P15 are pH3+ and G0/early G1 cells are negative for pH3.

5000 cells for each gated population were collected using 4-way purity using either a 85 or 100 µm nozzle, into 1.5 ml Eppendorf Protein Lo-Bind tubes. Four biological replicates were collected. An interphase library sample were collected by combining 300,000 cells of G0/G1, S, and G2 populations. A mitotic library sample was composed of 800,000 mitotic cells gated by high DNA content and high Histone H3 Ser28 phosphorylation. Samples were centrifuged and supernatant removed before storing at −20 °C.

### In-cell digest

Cell sorted library samples, and unstained unsorted TK6 cells, were resuspended in DPBS at 2 - 5 million cells per ml and incubated with 1 µl (25 - 29 U) benzonase (Millipore) at 37 °C for a minimum of 1 hr. Trypsin was added to approximately 1:25 w/w and in-cell digested at 37 °C for ∼16 hrs. Digests were acidified with TFA and desalted over Sep-Pak C18 cartridges (Waters) and dried.

Individual populations of 5,000 cells were diluted with 40 µl PBS and incubated with 0.25 µl [6 – 7 U] benzonase at 37 °C for a minimum of 1 hr, then digested with 50 ng trypsin (∼1:10 w/w) at 37 °C for ∼16 hrs. Samples were acidified with TFA and desalted over self-made C18 columns with 3 Empore C18 disks [10] and eluted directly into Axygen™ 96-well PCR Microplates (Fisher Scientific) and dried.

### High pH reverse phase fractionation

Approximately 100 µg interphase, mitotic, and unsorted TK6 cell digests were fractionated by high-pH reverse phase chromatography using an Ultimate 3000 HPLC (ThermoFisher Scientific) and a 1 x 100 mm 1.7 µm Acquity UPLC BEH C18 column (Waters). Peptides were separated using a constant 10 mM ammonium formate (pH 10) and a gradient of water and 100% acetonitrile. Peptides were loaded at 1% acetonitrile followed by separation by a 48 min multistep gradient of acetonitrile from 3% to 6%, 25%, 45% and 80% acetonitrile at 4, 34, 44, 45 minutes, respectively, followed by an 80% wash and re-equilibration. Fractions were collected at 30 sec intervals resulting in 96 fractions which were concatenated into 12, and 1 µg aliquots dried.

### LC-MS/MS

Peptide samples were resuspended in 0.1% TFA. Approximately 0.5 µg of library fractions were injected for DDA LCMS analysis. A volume equal to half the cell population (equivalent to ∼2,500 cells) was injected and analysed twice by AMPL to produce two technical replicates for each of the four biological replicates. An Ultimate 3000 RSLCnano HPLC (Dionex, Thermo Fisher Scientific) was coupled via electrospray ionisation to an Orbitrap Elite Hybrid Ion Trap-Orbitrap (Thermo Fisher Scientific).

Peptides were loaded directly onto a 75 μm x 50 cm PepMap-C18 EASY-Spray LC Column (Thermo Fisher Scientific) and eluted at 250 nl/min using 0.1% formic acid (Solvent A) and 80% acetonitrile/0.1% formic acid (Solvent B). Samples were eluted over 90 min stepped linear gradient from 1% to 30% B over 72 min, then to 45% B over 18 min. AMPL analyses included up to 5 MS1 microscans of 1E6 ions in the Orbitrap at 120k resolution and with a 250 ms maximum injection time. MS1 scans were acquired over 350-1700 m/z and a ‘lock mass’ of 445.120025 m/z was used. This was followed by 5 data-dependent MS2 CID events (5E3 target ion accumulation) in the ion trap at rapid resolution with a 2 Da isolation width, a normalised collision energy of 35, 50 ms maximum fill time, a requirement of a 10k precursor intensity, and a charge of 2+ or more. Precursors within 5 ppm were dynamically excluded for 40 sec. DDA analyses were as for AMPL but with a single MS1 microscan with a 75 ms maximum injection time, followed by 20 CID events in the ion trap.

Libraries were acquired as for DDA analyses or acquired with 10 data-dependent MS2 HCD events at 30 NCE of 5E4 ions in the Orbitrap at 15k resolution and a maximum fill time of 100 ms, with a precursor intensity required to be at least 50k. For the sample preparation comparisons shown in Fig. 2, a 240 min gradient was used (1% to 30% B for 210 min, then to 42% B over 30 min). MS data was acquired as for DDA analysis described above with the exception that MS1 spectra were acquired at 60k resolution and MS2 events were acquired only on 2+ and 3+ precursors.

### MS/MS data analysis

Data was processed using MaxQuant version 1.6.2.6 [15]. LC-MS/MS data was searched against the Human Reference Proteome from UniProt including splice-isoforms (accessed October 23^rd^, 2017), which contains 93,613 entries, allowing for 2 tryptic missed cleavages, allowing for variable methionine oxidation and protein N-terminal acetylation. Carbamidomethyl cysteine modification was allowed only for samples that were alkylated by iodoacetamide. The parameter “Individual peptide mass tolerance” was selected for variable precursor mass tolerances, with 0.5 Da or 20 ppm mass tolerances set for ion trap or orbitrap fragment ions, respectively. A target-decoy threshold of 1% was set for both PSM and protein false discovery rate. Match-between-runs was enabled with identification transfer within 0.5 mins and a retention time alignment within 20 min window. Matching was permitted from the library parameter group, and ‘from and to’ the unfractionated parameter group. The parameter “Require MS/MS for LFQ comparisons” was deselected, and second peptide search was enabled. Both modified and unmodified unique and razor peptides were used for quantification. ‘Evidence’ and ‘proteinGroups’ output files were used for subsequent analysis in R.

### Match-between-runs FDR filtering

A reference sample was generated by lysing TK6 cells in DPBS with 2% SDS and cOMPLETE protease inhibitors without EDTA (Roche, 1x concentration) at 70 °C, homogenised with a probe sonicator and treated with benzonase. Protein was reduced with 20 mM TCEP for 2 hr before alkylation with 20 mM iodoacetamide at ambient temperature in the dark for 1 hr. Protein was precipitated with 4 volumes cold acetone at −20 °C overnight, washed with 100% cold acetone and 90% cold ethanol. Protein pellet was air dried before resuspending in DPBS and digesting with 1:50 w/w trypsin for ∼16hrs. Peptides were acidified, desalted, aliquoted, and fractionated as previously described. For isopropylation, 50 µg peptides were resuspended in 200 µl 90% acetonitrile containing 0.1% formic acid before addition of 50 µl acetone containing 36 µg/µl NaBH_3_CN. The reaction was conducted at ambient temperature for ∼16 hrs before quenching with ammonium bicarbonate, drying off solvent and desalting peptides over C18. For dimethylation, 50 µg peptide was resuspended in 200 µl DPBS before addition of 0.32% formaldehyde and 50 mM NaBH_3_CN. The reaction was conducted at ambient temperature for ∼16 hrs before quenching with ammonium bicarbonate and desalting peptides over C18. 200 ng of unmodified, dimethylated, and isopropylated peptides were analysed by AMPL and DDA, and unmodified fractionated peptide samples were analysis by DDA, as previously described. LCMS data were searched using MaxQuant, as previously described. Note that dimethylation and isopropylation modifications were not specified in in the search parameters.

### Cell cycle proteomic data analysis

All subsequent data analysis on the protein intensity table, including the analysis of pseudoperiodicity, was performed using R (v. 3.5.0) within the RStudio integrated development environment. The R scripts are available as Supplementary Data 1. The list of validated APC/C substrates was obtained from the APC/C degron repository (http://slim.icr.ac.uk/apc/). Proteins that contain D box, KEN and ABBA SLIMs in the human proteome were found using SLiMsearch with default settings (Disorder score cut-off: 0.30, Flank length: 5). In order to remove slight variations in total protein amount in each sample, protein intensities were divided by total intensities per sample and multiplied by 10^6^ to obtain intensities in parts per million (ppm). There are four biological replicates analysed in technical duplicate. As described above, sample analysis was completely randomized in the second technical repeat. Each technical repeat (i.e. set of four biological replicates) are considered as one ‘pseudotimecourse’ with samples in each biological replicate arranged in order from P1 to P16. Each of the two pseudotimecourse was then independently subjected to a Fisher’s test for periodicity, as implemented in the ptest R library (v. 1.0-8). Fisher’s periodicity test p-values were corrected for multiple hypothesis testing using the q value method as implemented in the qvalue R library (2.15.0). Those proteins that showed q values <0.10 in both sets of biological replicates and oscillation frequencies of either 0.0625 (1/16) or 0.125 (1/8) were classified as pseudoperiodic.

For clustering, protein ppm values were averaged (mean) to produce a single pseudotimecourse for each protein. These average abundance profiles were scaled using the base R function scale and subjected to hierarchal clustering using the Ward minimum variance algorithm. The appropriate range for cluster number was identified as 3 - 6 clusters using the ‘elbow method’, which involves plotting within-cluster sum of squares versus number of clusters. Bifurcating leaves of the subsequent dendrogram were swapped in order to produce a heatmap that follows a logical, sequential order of peak abundance, i.e. cluster 1 with highest abundance in P0-P8 and cluster 5 with peak abundance in P3-P7, etc.

For PCA and cell cycle state classification, scaled pseudotimecourses were used. Cell cycle states were classified using the k-NN model as implemented in the class R library (v. 7.3-15) using k = 6, with k being the number of nearest neighbours for classification. Three biological replicates were used as the training set and the remaining replicate was used as a test set.

For the pairwise comparison of the proteomes of P17 with P1 and P16, t-tests were performed on ppm intensities. Uncorrected p-values were plotted against mean fold change in order to identify candidate proteins that were specifically changed in abundance in P17.

## Results

### The ‘in-cell digest’: direct protease digestion of fixed cells

Based on previous work [1,16], we hypothesized that formaldehyde-induced modifications were of low stoichiometry and crosslink reversal may not be required for proteome analysis. Consistent with this idea, deep proteome analysis comparing human epithelial MCF10A cells fixed with 2% formaldehyde for 10 min, fixed and treated with heating to reverse formaldehyde crosslinks, or not fixed (Fig. 1A and Supp. Fig. 1A) showed no significant differences in protein and peptide coverage (Figs. 1B-C). These proteomes were analysed to a depth of ∼53,600 peptides and ∼7,700 proteins. We next used the error-tolerant MS search to find peptides chemically modified by formaldehyde. The pattern and frequency of detected mass shifts are remarkably similar between control and fixed samples (Suppl. Fig. 1B). From these observations, we concluded that under these controlled and mild fixation conditions, the stoichiometry of crosslinking and chemical modification by formaldehyde is sufficiently low such that the non-detection of modified and crosslinked peptides is not detrimental for characterization of proteomes to a depth of at least 7,700 proteins.

**Figure 1.**
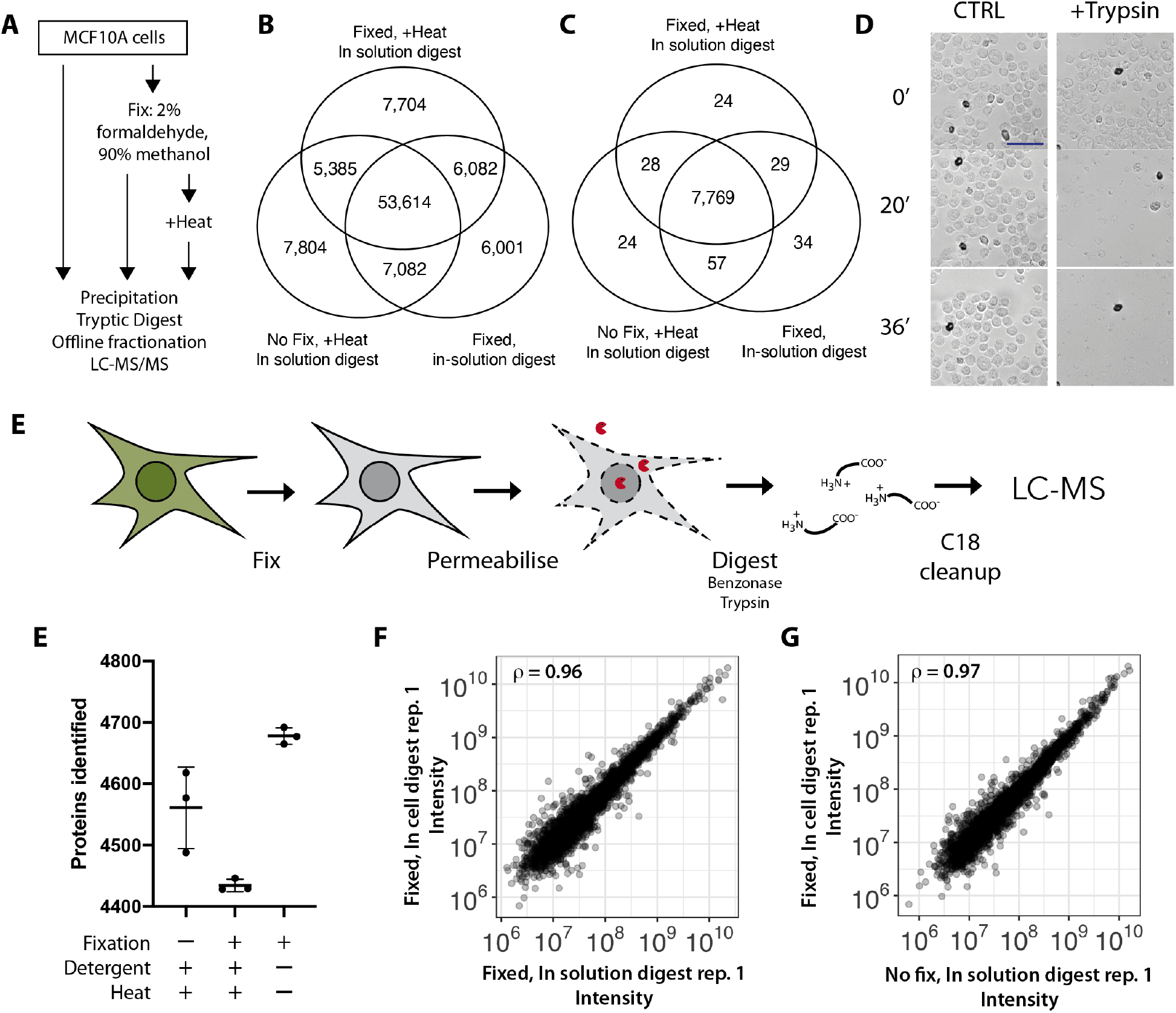
A simplified workflow for proteomics of formaldehyde fixed cells using in cell digestion. A) Proteomics workflow to assess impact of formaldehyde-induced modifications on proteome analysis. B, C) The number of peptides (B) and proteins (C) identified. D) Fixed and permeabilized cells treated either with DPBS (left) or with trypsin (right) were imaged at the indicated times in minutes. Scale bar is 50 µm. E) Schematic of the in-cell digest workflow. F-H) Comparison of the identification (F) and quantitative reproducibility (G, H) between in-solution and in-cell digests.

We next hypothesized that fixed cells may make suitable substrates for direct protease digestion. Digestion of fixed cells would significantly simplify the sample processing workflow by eliminating several steps, including detergent-based lysis, homogenization, heat treatment and detergent removal. We therefore treated fixed, permeabilized cells suspended in DPBS with either mock treatment (DPBS), or trypsin, and monitored cell morphology by brightfield microscopy. As shown in Fig. 1D, prominent structural features visible in control cells, such as the plasma membrane, nuclei and nucleoli, are degraded in a time-dependent manner by trypsin (see Supplementary Video 1). For LC-MS/MS analysis, fixed cells were also pre-incubated with benzonase to digest RNA and DNA oligonucleotides, which may interfere with downstream sample processing. The peptide-containing supernatant from the digest was then subjected to C18 purification prior to analysis by LC-MS/MS. As the digestion occurs within the fixed cells, we have called this approach an ‘in-cell digest’ (Fig. 1E).

As shown in Fig. 1F, the proteome coverages are similar for fixed cells processed by the in-cell digest method (∼4,678 proteins, n = 3), fixed samples that were subjected to the PRIMMUS protocol (∼4,446 proteins, n = 3) and extracts from non-fixed cells processed by precipitation (see Methods, ∼4,561 proteins, n = 3). We conclude that the proteome coverage from the in-cell digest is similar, or higher, than the other protocols tested.

We did not observe a broad bias in quantitation, as label free intensities measured in fixed cells prepared by the in-cell digest and by decrosslinking followed by an in-solution digest showed high correlation (Fig. 2G, ρ = 0.96). Similarly, a high correlation was observed between fixed cells prepared by the in-cell digest and non-fixed cells (Fig. 2H, ρ = 0.97). Few points lie off-diagonal, indicating that proteins showing a difference in intensity between methods are rare. We then tested if these off-diagonal proteins were enriched in any UniProt keywords or GO annotations using DAVID. The only terms that were significantly enriched in proteins showing lower intensity with the in-cell digest were associated with RNA-binding (FDR < 0.05, Suppl. Fig. 1C). Notably, these RNA-binding proteins are present in cells at high abundance. In contrast, proteins showing higher intensity with the in-cell digest are enriched in membrane proteins (FDR < 0.05, Suppl. Fig. 1D). Improved recovery of membrane proteins using the in-cell digest is consistent with previous results demonstrating that heat treatment can irreversibly precipitate membrane proteins.

**Figure 2.**
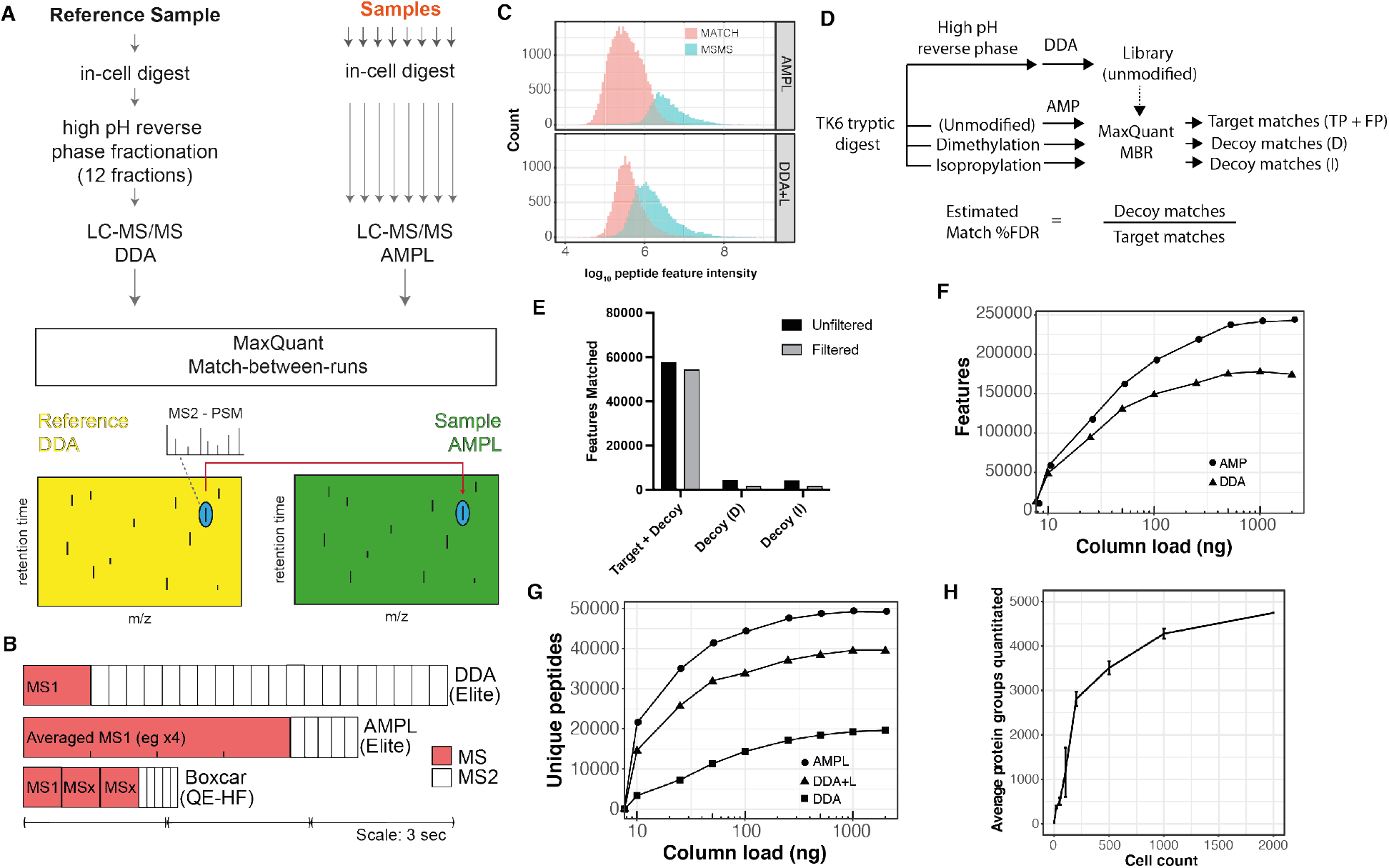
Averaged MS1 precursors with library matching (AMPL) increases peptide detection sensitivity. A) Schematic outlining the AMPL experimental design. B) Both the AMPL and BoxCar acquisition methods prioritise MS time to enhance MS1 scan quality. Schematic comparing duty cycles for DDA, AMPL and Boxcar acquisition methods on the indicated MS instruments (Orbitrap Elite, Orbitrap HF). The median peak width using our chromatographic setup with the Orbitrap Elite is ∼38 seconds. C) A comparison between AMPL and DDA+L, showing intensity distributions of peptide features identified by MS/MS (blue) and matching to identified library features (red). D) Schematic outlining experimental workflow for assessing match-between-runs FDR. E) Features matched in target and decoy proteomes before and after additional filtering based on match retention time difference, match m/z difference and match m/z error. F, G) Features (F) and unique peptides (G) detected in AMP(L) vs DDA. DDA+L is DDA with matching to a library. H) Proteome coverage versus cell number. The cell titration was performed in duplicate.

We conclude that the measurements of protein abundance from the in-cell digest are quantitative, reproducible and broadly comparable to conventional sample preparation methods. We note that each sample preparation method will have its own specific biases. In the case of the in-cell digest, the increased abundance of membrane proteins may more accurately reflect the abundance of these proteins in cells.

### Averaged MS1 Precursors with Library matching (AMPL) improves feature detection

To increase the sensitivity and detection speed of the Orbitrap Elite MS instrument, we utilised MS1-based identification and quantitation using accurate mass and retention time matching, as proposed originally by the Smith lab [18]. This approach has been recently demonstrated to be highly sensitive in an implementation called BoxCar [19]. The BoxCar method increases the signal-to-noise of trap-based MS by collecting ions using segmented, spaced windows. Peptide identification relies on MS1 feature matching to a reference library generated from a fractionated reference sample using the MaxQuant function ‘Match-between-runs’ (MBR). The library is analysed separately using data-dependent acquisition (DDA) and peptides are identified by MS2 and database searches.

As the BoxCar method cannot be directly implemented on the Orbitrap Elite, we developed a different approach to increase the dynamic range of MS1 feature detection. MS1 spectral averaging is frequently performed in direct infusion MS, but rarely employed in LC-MS bottom-up proteomics. We surmised that averaging several MS1 scans would improve signal-to-noise (S/N) and would rapidly plateau as it is known that averaging improves S/N by a factor of sqrt(n) where n is the number of spectra averaged. Features would then be matched between the single shot analyses to a fractionated reference library (Fig. 2A). We call this method Averaged MS1 Precursors with Library matching (AMPL), or AMP if no library is used. Like BoxCar, AMP(L) prioritises MS1 scans over MS2 scans as compared with DDA (Fig. 2B) and includes top-5 DDA MS/MS scans to ensure identification of features for accurate retention time alignment throughout the chromatographic separation.

We therefore tested AMPL by analysing 1µg on-column loads of MCF10A tryptic digests. A comparison of different MS1 scans (n = 1, 3, 4, 5) showed that the number of features and peptides identified saturates at n = 4 (Suppl. Figs. 2A-B). AMPL (n = 4) detects ∼278,205 features, representing a 20% increase compared to a standard Top 20 DDA acquisition using the same gradient (188,928 features). We reasoned that the additional peptides detected by AMPL originate from low-abundance features detected by virtue of the S/N increase due to averaging. Fig. 2C compares the peptide intensity distributions between DDA-L and AMPL. The distributions are bimodal, with MS/MS-dependent identification biased towards higher intensity features (cyan). Consistent with the idea that AMPL improves S/N, AMPL detects a higher number of matched features (pink) in the low abundance regime. Similar to previous MS1-based matching approaches, AMPL shows higher data completeness (4,411 proteins with intensities measured in all 10 replicates) as compared with DDA-L (3,493 proteins) and DDA (2,865 proteins) (Suppl. Fig. 2C).

MS1-based matching significantly increases the sensitivity, coverage and data completeness of MS-based proteomics. However, the lack of MS2-based identification for these matched sequences could potentially increase the false discovery rate (FDR). We estimated the matching FDR by using an empirical ‘target-decoy’ approach where decoy proteomes created by chemical modification (dimethylation and isopropylation) are matched against an unmodified library (Fig. 2D). Whereas matches to the target proteome will contain both true and false positives, matches to the decoy proteomes should contain exclusively false positives (with the rare exception of peptides containing an N-terminal acetyl group a C-terminal arginine, which are not dimethylated/isopropylated). ∼32% of the features are assigned a peptide sequence when the target, unmodified proteome is matched against an unmodified library (Suppl. Fig. 2D). By contrast, only ∼2% of the features are matched in the decoy samples (Suppl. Fig. 2D). Using this approach, the estimated match FDR is 7.4%. To reduce the FDR <5%, we applied more stringent thresholds for match time, match m/z and match m/z error (2.5 σ for match time, 3 σ for match m/z and match m/z error, Suppl. Fig. 2E-F). Application of these thresholds reduced the estimated FDR to 3.1-3.4% (Fig. 2E) while retaining 96% of the matches in the target dataset.

The improvements in detecting low abundance features suggest that AMPL may be well suited to analysis of low sample loads. AMP (i.e. no library) consistently detects more features than DDA (Fig. 2F), which leads to significant improvements in peptide coverage (Fig. 2G). For example, at 10 ng loading, 21,483 unique peptides are quantitated by AMPL versus 14,702 by DDA-L, representing a 46% increase in coverage. AMPL provides 150-535% improvement relative to conventional DDA with no library and 24-46% improvement relative to DDA-L for protein coverage at all tested column loads with greatest gains observed at low column load.

As shown in Fig. 4G, AMPL detects a slightly higher number of peptides in 10 ng on-column load as DDA with 1 µg load, demonstrating a >100x increase in sensitivity. A 10 ng on-column load is equivalent to the protein content of ∼67 cells based on the protein per cell measured in bulk assays. However, the effective number of cells required for proteome analysis is frequently much higher. This is due to losses during sample preparation. We reasoned that these losses are significantly reduced using the streamlined in-cell digest.

We combined the in-cell digest with AMPL to analyse FACS collected TK6 cells, a human lymphoblastoid cell line with a stable near-diploid karyotype. Notably, TK6 cells are smaller than typical adherent human cell lines, such as HeLa and MCF10A. Being cultured in suspension, TK6 cells are amenable towards cell separation techniques, including fluorescence-activated cell sorting (FACS) and centrifugal elutriation, without requiring cell dissociation, which can induce physiological perturbations.

Fig. 2H shows the result of a cell titration analysis of S-phase cells performed in duplicate whereby two aliquots at each indicated cell number (2,000 cells to 0 cells) were collected by FACS from the same starting cell population. ∼4,500 proteins were quantitated with 2,000 cells, with 4,480 proteins reproducibly quantitated in two technical repeats. At the lower end of the cell titration, 2,933 proteins on average were quantitated from 200 cells. We note that below this number of cells, we observe a higher variability in proteome coverage, which will need to be addressed by further optimization. Indeed, while approx. 30 proteins were detected in single cells, with 17 reproducibly detected, nearly all of these proteins were also detected in the background samples (‘0 cells’).

We conclude that combining in-cell digest and AMPL enables characterization of proteomes of 2,000 cells to a protein depth comparable to conventional single shot DDA analysis of 1 µg on-column loads. The advanced PRIMMUS method presented here significantly reduces the number of cells required, i.e. ∼10^3^ versus ∼10^5^ with low estimated match FDR (<3.5%).

### High temporal resolution analysis of an unperturbed cell cycle using PRIMMUS

The process of normal cell division requires linear progression through several cellular states i.e. S- and M-phases) in which DNA replication and mitosis must occur in sequential order. These states can be further resolved. DNA replication, for example, occurs in a temporally and spatially patterned manner, with euchromatic genomic regions replication first before heterochromatin-dense regions. Similarly, M-phase can be resolved into prophase, prometaphase, metaphase, anaphase I, anaphase II and telophase based on cytological features. Some of these phases, including telophase are exceptionally rare in asynchronous cells and are not amenable for collection by FACS in numbers required for typical proteomic analysis. We therefore developed an advanced PRIMMUS workflow incorporating the in-cell digest to target these rare cell states and carry out a high temporal resolution analysis of proteome variation across 16 cell cycle subpopulations, including 8 interphase and 8 mitotic states. This fractionation-based approach to separating cell cycle phases relies on continuous cell trajectories, such as cell cycle progression in asynchronous populations that are unperturbed by drug-based synchronisation.

TK6 cells were immunostained for DNA content, cyclin B, cyclin A and histone H3 phosphorylation (Ser28), which are all markers of cell cycle progression. Cells were then separated into 16 cell cycle populations (P1 – P16) (see Suppl. Fig. 3 for the full gating strategy). Biochemical differences are used as a surrogate for time and cell cycle progression. Based on past literature [20,21] and our previous data [1], we have correlated these biochemical changes with specific cell cycle states (as illustrated in Fig. 3B, top). For example, cyclin A and cyclin B levels are used to discriminate mitotic subphases, as they are degraded during prometaphase and the metaphase-to-anaphase transition, respectively.

**Figure 3.**
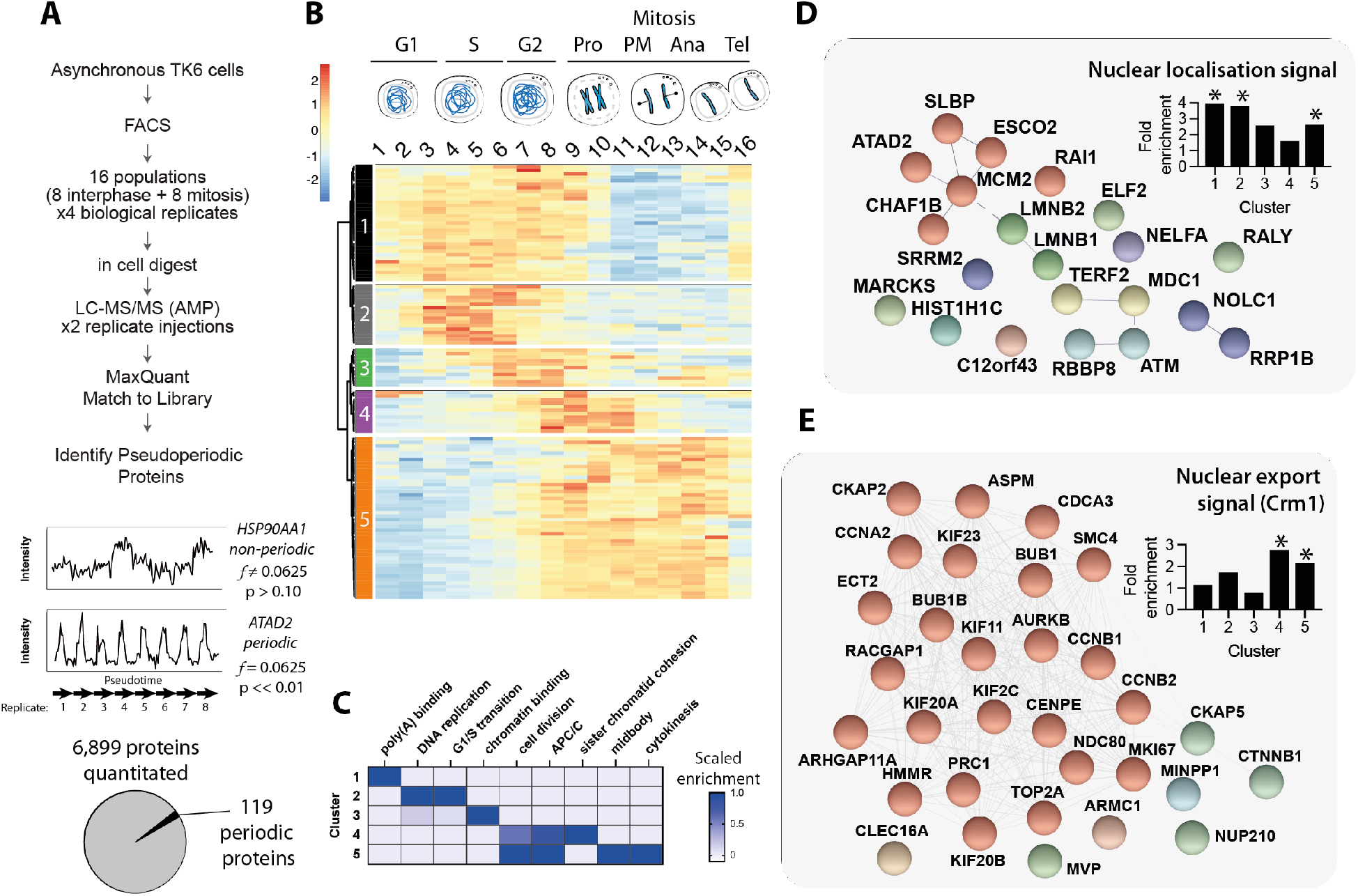
High resolution proteomic analysis of an unperturbed cell cycle. A) Schematic describing the experimental design and workflow. Details for protein abundance normalization and pseudoperiodicity analysis can be found in the Experimental methods section. B) Heatmap of the 119 identified pseudoperiodic proteins (PsPs) organized by cluster. Schematic above heatmap shows indicative cell cycle stages isolated by FACS (full gating strategy shown in Suppl. Fig. 3). C) Enriched UniProt keywords by cluster. D) Proteins in clusters 1 and 2 containing a putative nuclear localization signal (NLS). E) Proteins in clusters 4 and 5 containing a putative nuclear export signal (NES). * indicate p < 0.01 (Fisher’s exact test).

**Figure 4.**
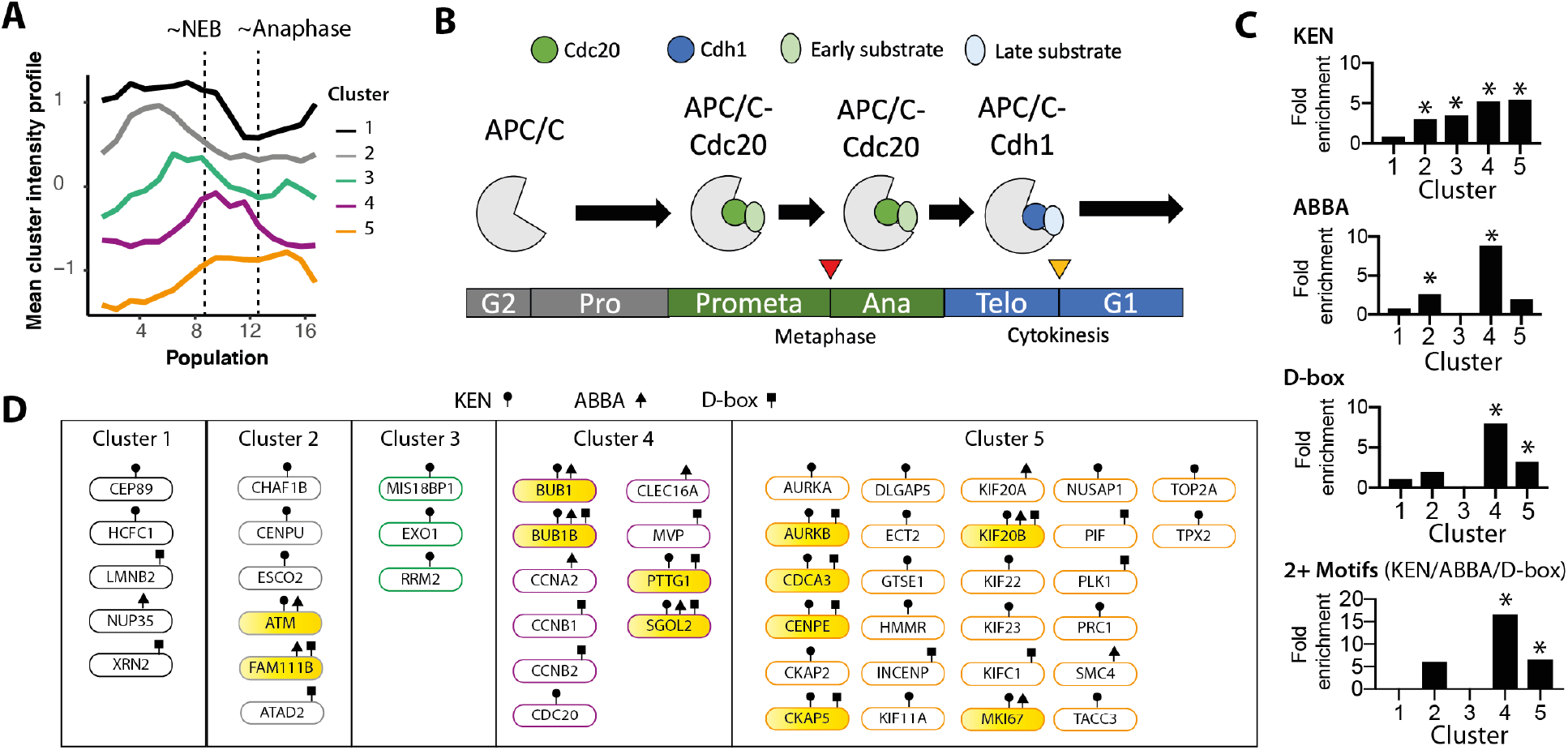
Characterisation of potential APC/C substrates. A) Mean intensity profile for five clusters shown in Fig. 3B. B) Schematic illustrating regulation of APC/C substrate choice switch during mitosis by co-activators Cdc20 and Cdh1. C) Enrichment analysis of SLiMs that control interaction with the APC/C. D) Proteins with at least one APC/C SLiM grouped by cluster. The yellow fill indicates proteins that contain 2 or more SLiMs.

The rarest target population are cells in late anaphase of mitosis, which are present in 0.01% of an asynchronous TK6 culture. Proteome characterisation of these cells, previously challenging due to lack of sensitivity, is now possible with the in-cell digest. Four separate cultures of TK6 cells were independently FACS separated into 16 populations. For each population, 5,000 cells were collected and processed using the in-cell digest. Collection of 5,000 cells provided sufficient material for duplicate injections for LC-MS/MS analysis by AMPL with DDA libraries generated from interphase, mitotic and asynchronous cells. The data were then processed by MaxQuant with MBR and filtered by match parameters as discussed above to generate a dataset with 6,899 quantitated proteins (Suppl. Table 1).

Next, to identify cell cycle regulated proteins, we treated each set of 16 populations as an ordered series of related biochemical states. These states were projected onto a temporal axis (i.e. cell cycle progression). A single replicate series of ordered cell states constitutes a ‘pseudotimecourse’ (Fig. 3A, bottom). We then applied the Fisher’s periodicity test to identify ‘*ps*eudoperiodic *p*roteins’ (PsPs), i.e. proteins abundance patterns that showed periodic behavior across the four pseudoperiodic timecourses. In order to increase robustness, the periodicity test was separately performed on each technical repeat and only those proteins showing periodicity in both were designated as PsPs. Fig. 3A (bottom) shows the abundance profiles for heat shock protein HSP90AA1 and ATPase AAA domain-containing protein ATAD2 as an example non-PsP and PsP, respectively. ATAD2 shows highly reproducible abundance variation in all 8 pseudotimecourses, with peak abundance in S-phase populations (P5-P6), consistent with previous reports [22]. In total, 119 PsPs were identified using these criteria (Fig. 3A, bottom, Suppl. Table 2).

Hierarchal clustering of the 119 PsPs identified five major classes of protein abundance patterns (Fig. 3B). Each cluster shows peak abundance at different times during cell cycle progression. The gene ontology terms enriched for each cluster reflects key processes and/or compartments associated with that period during cell cycle progression. We also assessed enrichment in short linear (sequence) motifs (SLIMs). SLIMs mediate protein-protein interactions that lead to changes in post-translational modification, stability and/or subcellular localization of a protein. Using the eukaryotic linear motif (ELM) database [23], we identified SLIMs that are enriched in each cluster (p < 0.01, Fisher’s exact test, Suppl. Table 3).

Cluster 1 proteins show high abundance in interphase, which decreases in early mitosis (P8-P10) and recovers slightly in late mitotic populations (P15-P16). This cluster is highly enriched in proteins with a monopartite nuclear import signal sequence (Fig. 3D) and in contrast to other clusters, do not show any enrichment for the Crm1-mediated nuclear export signal sequence (Fig. 3E). Most proteins in this cluster are either RNA- or DNA-binding (26 / 33). For example, several mRNA splicing factors are in this group, including serine/arginine-rich proteins (SRRM2, SRSF2, SRSF3, SRSF5, SRSF6, SRSF10/TRAB). These proteins reproducibly decrease in abundance in mitosis, but with a small fold change (< 2-fold) than key cell cycle regulators, e.g. cyclin B1 (> 4-fold). The stability of the SR proteins is regulated by nucleocytoplasmic shuttling. For example, SRSF1 is stable in the nucleus but has a short half-life in the cytoplasm [24]. Proteasome-dependent degradation of SR proteins is dependent on the RS domain, which is shared among SR proteins [25]. Cluster 1 is also enriched in poly(A)-binding proteins in the nucleus that are involved in pre-mRNA and ribosomal RNA processing, e.g. XRN2, NOLC1. The remaining proteins with no known or anticipated oligonucleotide-binding properties are enriched in cytoskeleton-binding factors, e.g. the actin-binding protein MARCKS, CCDC6, CEP89 and DBNL.

Cluster 2 proteins peak in late G1/S. Nearly all proteins in this cluster are directly involved in DNA replication, establishment of nascent chromatin, or the G1/S transition (Fig. 3C). In this cluster are three members of the MCM helicase (MCM2, MCM5, MCM6), the replication-dependent histone chaperone (CHAF1B) and the histone mRNA stem-loop binding factor SLBP, which is essential for the synthesis of histones for incorporation into newly synthesized DNA in S-phase. This cluster also includes the DNA damage checkpoint kinase ATM, which is important in resolving endogenous replication stress [26].

Cluster 3 shows peak abundance in late S, G2 (P6 to P8) and decreased abundance in early-mid mitosis (P9 to P11). Three proteins show >5-fold decrease in abundance by mid-mitosis with low or undetectable levels in late mitosis: GMNN, RRM2 and PAF/KIAA0101. All three are targeted for degradation in late mitosis and G1 by the APC/C-Cdh1. The remaining proteins in the cluster show an increase in S/G2 phase and a decrease in prophase/prometaphase (P9-P12), followed by a slight recovery in abundance in late mitosis. These include sororin/CDCA5, which functions in sister chromatid cohesion establishment, and MIS18BP1, which facilitates loading of the centromere-specific histone in late mitosis and G1. This cluster is enriched in chromatin-binding factors, including TRIM28/KAP1, EXO1, sororin, PAF and MIS18BP1.

Clusters 4 and 5 show peak abundance during mitosis and contain the largest proportion of proteins with either known direct roles in mitotic progression or targeted for degradation in mitosis (9/12 for cluster 4, 38/46 for cluster 5). The feature that distinguishes clusters 4 and 5 is the mitotic abundance pattern. Cluster 4 proteins show decreased abundance in earlier mitotic populations, particularly in P11 – P12, coincident with the onset of cyclin A2 and cyclin B1 degradation (c.f. Fig. 5B). The three mitotic cyclins detected (A2, B1 and B2), the spindle assembly checkpoint kinases BUB1 (Fig. 6F) and BUB1B (BubR1), the kinesin-8 family member KIF18B, securin (PTTG1) and shugoshin (SGO2) are in this cluster. Functionally, this cluster is characterized by proteins that protect sister chromatid cohesion (securin, shugoshin) and the spindle assembly checkpoint that prevents anaphase (Bub kinases) while proper microtubule attachment and biorientation of chromosomes takes place.

**Figure 5.**
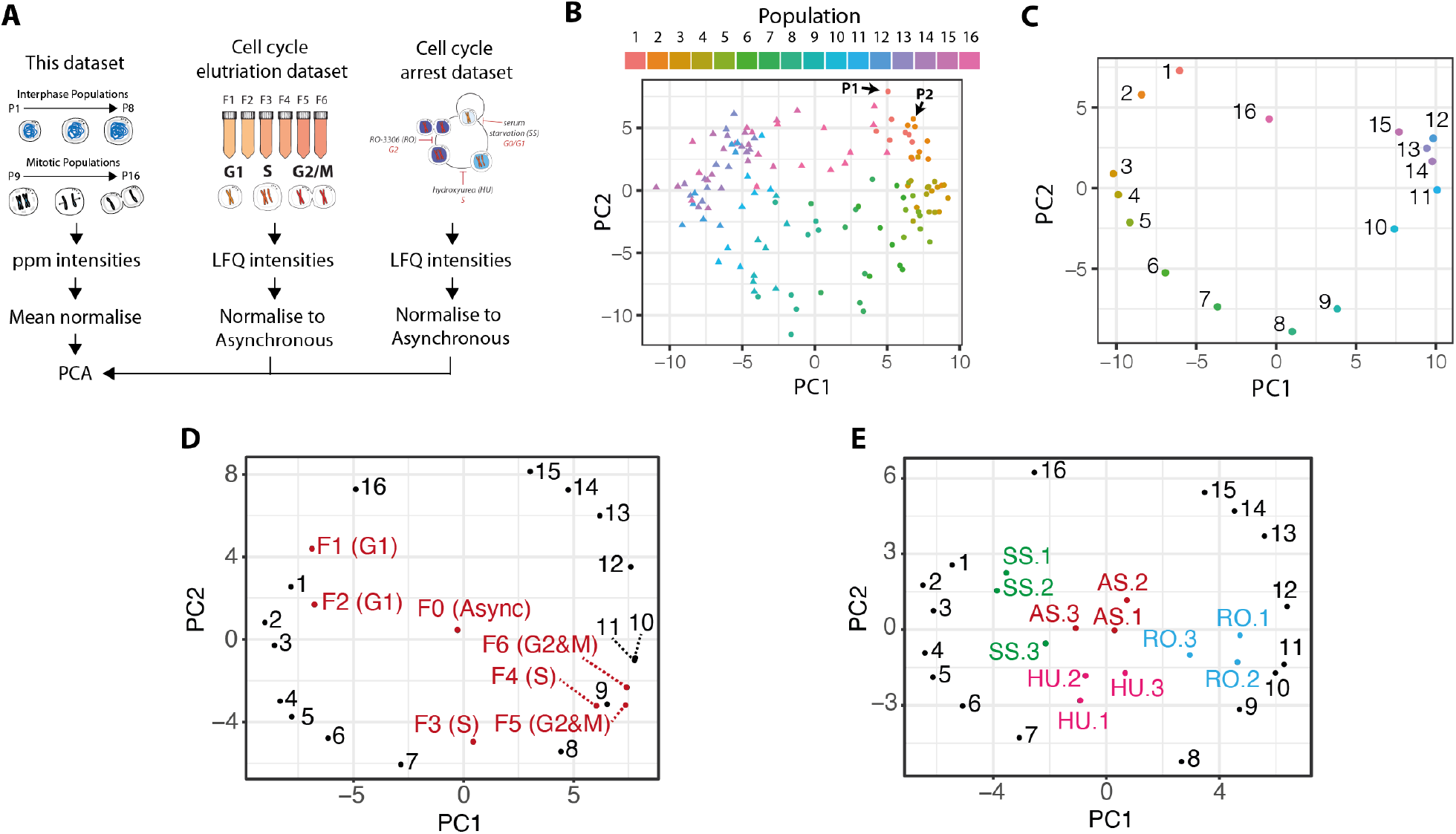
The 119 PsPs form a signature that classifies cell cycle states. A) Schematic illustrating the approach to compare the signature across datasets. B, C) PCA of the 16 cell cycle populations in this study using the 119 PsPs as features using either individual replicates (B), or mean abundances (C). D, E) PCA as in (C) with samples from elutriated (D) or from cell cycle arrest (E) label free datasets using NB4 cells.

**Figure 6.**
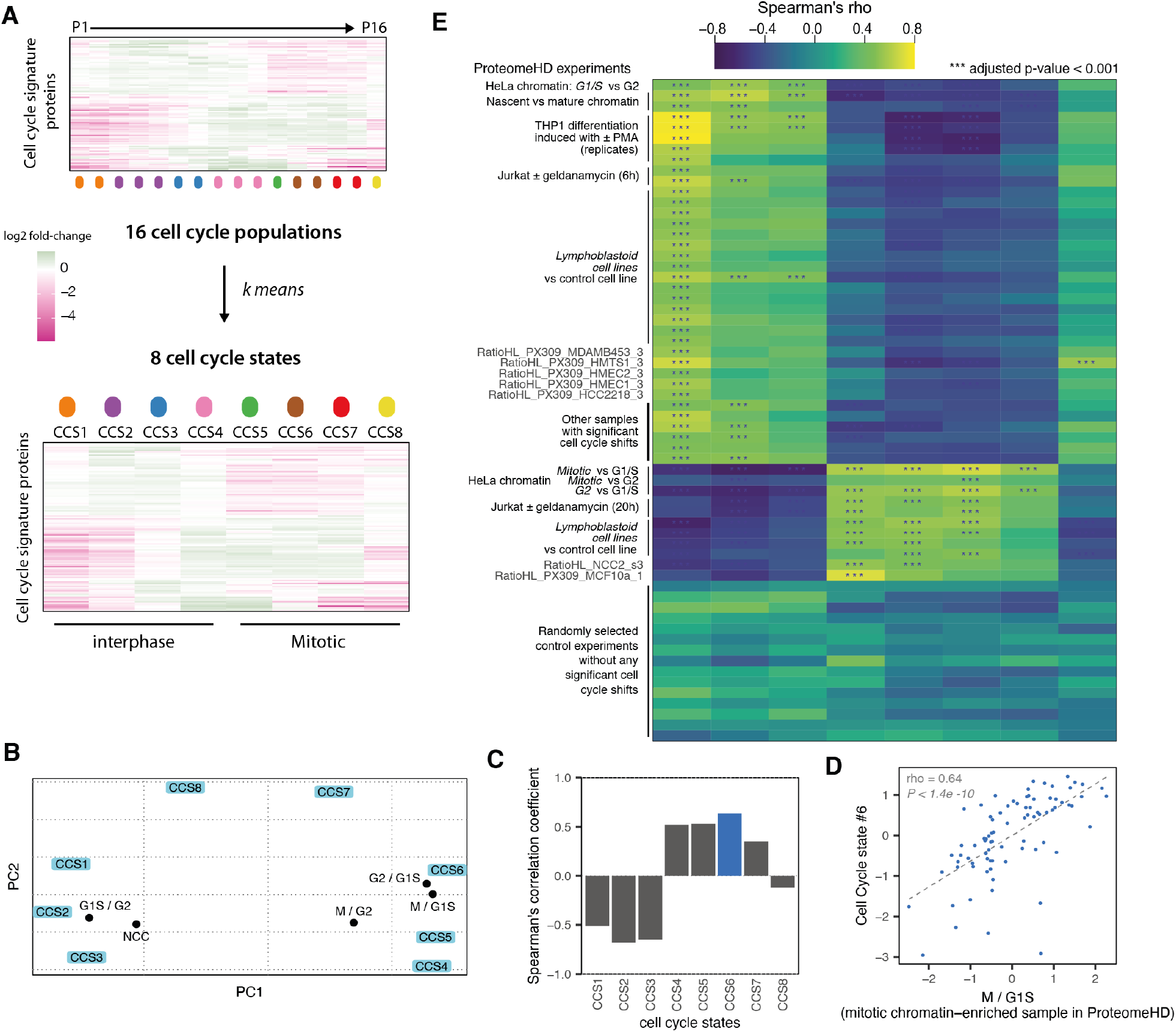
Unbiased assignment of cell cycle states across hundreds of proteomes in ProteomeHD. A) Schematic of how the 16 cell cycle populations were aggregated into 8 cell cycle states (CCS). B) Assignment of chromatin proteomes from nascent chromatin capture (NCC) and chromatin enrichment proteomics (ChEP) SILAC experiments. NCC samples were enriched in S-phase. G1S, G2 and M samples were treated with thymidine, RO-3306, and nocodazole, respectively. C) Spearman rank correlation coefficient for the 8 cell cycle states for M / G1S. This sample has the highest correlation with CCS6 (blue fill). D) Correlation scatterplot between CCS6 and M / G1S. E) Heatmap showing Spearman correlation coefficients for 47 experiments (out of 294) that show an enriched CCS and 15 randomly selected experiments that have no CCS enrichment.

By contrast, cluster 5 proteins show a significant increase in abundance at the end of interphase (P7 – P8) with peak abundance throughout mitosis (P9 – P15) and a significant decrease only in the last population (P16), i.e. cells undergoing mitotic exit. Example proteins include the catalytic E2 subunits of the APC/C (UBE2C, UBE2S), the chromosome passenger complex (AURKB, INCENP, BIRC5 – Survivin, CDCA8 – Borealin), polo kinase (PLK1) and the spindle-associated protein FAM83D. Both aurora kinases (Aurora A and Aurora B) are known to relocalise to the central spindle after anaphase onset. Aurora B activity is crucial for cytokinesis, the final step in cell division.

Clusters 4 and 5 are strongly enriched in the Crm1-mediated nuclear export signal (NES, Fig. 3E). 8/12 proteins in cluster 4 match the NES consensus. Notably, cluster 4 includes cyclins B1 and B2, and constitutive export of cyclin B-CDK from the nucleus is important in preventing premature mitotic entry. Exclusion from the nucleus of other proteins within these two clusters (Fig. 3E) may also be important in preventing premature activation of processes that are normally restricted to mitosis.

We identified PsPs that have no reported function in cell cycle control. These novel cell cycle regulated proteins may, like many of the other proteins identified in this manner, have significant roles in cell cycle progression. These candidates include EXO1, the DNA helicase PIF1, the guanine-exchange factor NET1 and the serine protease FAM111B.

### Analysis of mitotic protein abundance dynamics in unperturbed cells

A major difference between the clusters is the timing of protein abundance decrease (Fig. 4A). A critical regulator of protein abundance during the cell cycle is the anaphase promoting complex/cyclosome (APC/C). The APC/C is an E3 ubiquitin ligase and is active during the mitotic and G0/G1 phases of the cell cycle [27,28]. Its substrates include key regulators of the cell cycle, including cyclin A2 and cyclin B1. Ubiquitination of APC/C substrates is tightly temporally controlled, with APC/C substrate specificity changing during the cell cycle (Fig. 4B). This is mediated through changes in the APC/C co-activators and substrate recognition factors, Cdc20 and Cdh1. While APC/C-Cdc20 is active in early mitosis, the substrate receptor changes to Cdh1 in late mitosis, thereby conferring a temporal order to substrate degradation. Cdc20 is itself a substrate of the APC/C-Cdh1, allowing for switch-like handover in substrate receptor control.

25 PsPs (out of 119) are experimentally validated APC/C substrates [29], and of these, 24 are found in clusters 3, 4 and 5. Substrate recognition by APC/C-Cdc20 and APC/C-Cdh1 is mediated by the interaction between WD40 domains on the APC/C- (Cdc20/Cdh1) and SLIMs found on substrates. The KEN and D-box (RxxL) degrons are well documented SLIMs that bind both APC/C-Cdc20 and APC/C-Cdh1, with APC/C-Cdh1 having a preference for the KEN degron. A third SLIM called the ABBA motif was shown to be important in substrate recognition by APC/C-Cdc20 [30].

Fig. 4C shows the enrichment profile of these SLIMs across the six clusters. The KEN motif is comparably enriched in 4 out of the 5 clusters (Fig. 4C, top), with highest enrichments for the mitotic phase-peaking clusters (clusters 4 and 5). The frequencies range from 25% of the proteins in a cluster having the KEN motif (cluster 2) to 43% (cluster 5), representing a 3-5-fold enrichment over the background frequency (8%). All four clusters show low to non-detectable abundance in P16, P1 and P2, i.e. mitotic exit and G0/early G1 when APC/C-Cdh1 is active. In total, 35 cell cycle regulated proteins contain a KEN SLIM, approximately 50% (18 proteins) that have been experimentally characterized as APC/C substrates. The remaining uncharacterized 17 proteins are excellent candidates to be APC/C-Cdh1 substrates. Consistent with this prediction, cluster 1, which is the only cluster showing no enrichment for the KEN motif, contains proteins that have on average, higher abundance in G0/early G1.

Six out of 12 proteins that peak in mid-mitosis (cluster 4) contain the RxxL D-box sequence. The 50% frequency is ∼8-fold higher than the background frequency (6%). By contrast, the fold-enrichment is considerably lower in the other clusters (Fig. 4C). Similarly, 5 out of 12 proteins contain the ABBA motif (42%, Fig. 4C), representing a ∼9-fold enrichment over the background frequency (5%). D-box and ABBA motif-containing proteins in this cluster are mostly mutually exclusive (Fig. 4D). Of the D-box and ABBA motif containing proteins, two have not been previously experimentally characterized as APC/C substrates: MVP and CLEC16A.

Cluster 4 is highly enriched in proteins containing more than one SLiM (KEN/D-box/ABBA, Fig. 4C, bottom) and two proteins in this cluster contain all three SLIMs: BubR1 (BUB1B) and shugoshin-2 (SGOL2). KIF20B is the only other PP that has all three SLiMs and is in cluster 5. BubR1 has been demonstrated to interact with APC/C through these three SLiMs and acts as a pseudosubstrate to inhibit APC/C activities in spatiotemporally controlled manner [31]. It would be interesting to test the role of these SLiMs in the other two proteins (SGOL2 and KIF20B). For example, SGOL2 is important in protecting sister chromatid cohesion and the spindle assembly checkpoint [32].

### Proteomic assignment of cell cycle states

MS-based single cell proteome analysis is an emerging area. Recent advances in miniaturized sample preparation [11],[5,9,10] suggest that routine proteome analysis of single somatic mammalian cells will be possible in the near future. In comparison, single cell transcriptomics as a mature field with commercial kits now available. In single cell RNA-seq (sc-RNAseq) analysis [33], the deconvolution of cell cycle state has been critical [34,35] This is because cell cycle variation contributes significantly to the variation observed in a cell population. For example, to identify cell fate trajectories during differentiation, researchers relied on reference cell cycle regulated genes in order to identify the effect of cell cycle variation in the gene expression differences observed [36]. A validated reference set of cell cycle regulated proteins will be important for the biological interpretation of single cell proteomic datasets.

We tested whether the abundances of the PsPs determined in this study were sufficient to assign specific cell cycle states to cellular proteomes (Fig. 5A). The abundance patterns for the 119 proteins for each sample (16 timepoints x 8 replicates = 128 samples) were subjected to principal component analysis (PCA). The two major principal components, PC1 and PC2, explain 53% and 20.5% of the variance, respectively, as shown in Fig. 5B. Interphase (circles) and mitotic (triangle) are separated predominantly along PC1. To a lesser extent, subphases within each (for example, see arrows indicating P1 and P2) are separated along both principal components. Moving counterclockwise, starting from the top right for P1, the samples clearly follow a trajectory that reflects the position of each sample in the cell cycle, starting from early G1 (P1 and P2) to mitosis (left side, triangles). Telophase/cytokinesis populations (P16, pink triangles) are situated between the other mitotic populations and P1. To ease visualization, the PCA was repeated using mean values per population (Fig. 5C). Using unbiased, unsupervised methods, the PCA has arranged the populations into a cell cycle ‘wheel’, suggesting a largely continuous process with the major separation along PC1 correlated with interphase (P1 - P8) versus mitosis (P9 - P16). It is less clear what is the major correlate for PC2. We note however that there is a correlation with APC/C activity, with active APC/C in populations with positive values along PC2 (early G1 and end of mitosis) and inactive APC/C in populations with negative values (S and G2).

Detection of relevant features is essential as PCA analysis of the entire proteome dataset does not result in cell cycle separation. Repeating the PCA analysis with cyclin A2 and cyclin B1 removed essentially produces identical results, which indicates that the relationships produced by using ∼119 cell cycle marker proteins are robust towards the absence of individual proteins, including key proteins that drive cell cycle progression. This robustness will be important in assigning cell cycle states in diverse datasets, as described below.

We then asked whether the PCA classification could be used to assign cell cycle states to cellular proteomes obtained in published cell cycle fractionation and arrest experiments. Human promyelocytic leukemia cells (NB4) were fractionated by centrifugal elutriation into different cell cycle populations (Fig. 5A, middle) [37]. There are seven fractions (F0 – F6), which correspond to asynchronous (F0), and samples enriched in G1 (F1 – F2), S (F3 – F4), and G2&M (F5 and F6). In a separate experiment, NB4 cells were arrested in G0-phase, S-phase, and G2 phase, respectively, using serum starvation (SS), hydroxyurea (HU), and the drug RO-3306 (RO) [2]. Label free quantitation (LFQ) intensities were normalised to asynchronous cells and these ratios were combined with mean-normalised data from this dataset prior to PCA.

Figs. 5D and E shows the combined PCA plots for the elutriation and arrest datasets, respectively. The NB4 cell populations are broadly separated according to the appropriate cell cycle phase. For example, as shown in Fig. 5D, F1 and F2 are positioned nearby P1 (early G1). F3 is in-between P7 and P8 (late S/G2) and F4 is near P9 (late G2/early mitosis). F5 is closest to P9 whereas F6 is in-between P9 and P10 (late G2/early mitosis). The nearest population for F1 and F2 is P1, which is an early G1 population. F3 is in-between P7 and P8, which are late S/G2 populations. In Fig. 5E, the SS samples are nearest the early G1 populations, P1-P4. The HU samples are in-between P7 and P8, which are late S/G2 populations. The RO samples are positioned near P9 – P11, which are late G2/early mitotic populations. We conclude from these data that this signature can be used to classify cell cycle enriched label free proteomes.

We next tested if the cell cycle signature can be broadly applicable to assign cell cycle state to a proteome. To do this, we made use of a large set of stable isotope labelling by amino acids in cell culture (SILAC) datasets curated in proteomeHD [38]. Incomplete synchrony and/or cell cycle enrichment will generally lead to much poorer purities compared to FACS. This lowers the resolution of classification for bulk population samples, which likely contain mixtures of different phases unless purified by FACS. This will not be the case for single cell proteomes, which will be by definition in a single cell state.

To facilitate assignment of cell cycle states to partially or completely asynchronous bulk populations, we first used k-means clustering to reduce the number of classes from 16 populations to 8 cell cycle states (CCS) (Fig. 6A, Suppl. Table 4). PCA using these 8 CCS also shows the cell cycle ‘wheel’ (Fig. 6B). We then mapped chromatin proteomes (nascent chromatin capture, NCC and chromatin enrichment proteomics, ChEP) from synchronized cells, arrested with thymidine (G1/S), 3 hr thymidine release (NCC), RO-3306 (G2), or nocodazole (M) (Fig. 6B). Although these samples were from a different cell type than our cell cycle signature data (HeLa vs TK6), and had been processed differently (chromatin-enriched vs in-cell digest) as well as quantitated differently (SILAC vs label-free), these samples group according to the appropriate cell cycle phase. For example, the G2 and M-phase samples are grouped between CCS6 and CCS5, which are early-to-mid mitotic states. By contrast, G1/S and NCC samples are grouped with CCS2 and CCS3, which are G1/S states.

One challenge for the systematic classification of a heterogeneous set of proteomics data are missing values, because not all of our 119 signature proteins were detected in all experiments in ProteomeHD. We therefore employed Spearman rank correlation to correlate the abundance of the signature proteins in these chromatin proteomes with the 8 CCS. For example, the M/G1S proteome shows the highest correlation with CCS6 (Figs. 6C-D), which is a mitotic state.

We subsequently applied this correlation approach systematically to all 294 experiments in ProteomeHD. We found that∼15% of the experiments in ProteomeHD (47 out of 294) showed a high and significant correlation with one or more CCS (Suppl. Table 5). Many of these experiments involve a cell cycle perturbation, including the NCC and ChEP experiments described above (Figs. 6B-D). These experiments also include other types of perturbations, including differentiation, where cell cycle arrest is an expected direct consequence. For example, proteomes from THP-1 monocytic cells treated with phorbol myristate acetate (PMA) ester are highly correlated with G1 cell cycle states. PMA treatment induces terminal differentiation of these cells and leads to cessation of cell proliferation. In total, ∼50% of the proteomeHD experiments highly correlated with a CCS can be linked directly to cell cycle arrest.

From these data, we conclude that the signature robustly and accurately assigns CCS across far ranging experimental contexts, cell types and quantitation strategies.

The remaining experiments with high correlation have less obvious links to cell cycle. For example, Jurkat T cells are treated with the HSP90 inhibitor geldanamycin for either 6 h or 20 h (Fig. 6E). Proteomes from 6 h treatment are highly correlated with CCS1 (early G1). By contrast, proteomes from 20 h treatment are highly correlated with the G2/mitotic states, CCS4 and CCS6. Geldanamycin has been reported to arrest cells in G1 or G2 phases of the cell cycle. Interestingly, flow cytometry analysis of cells treated with geldanamycin for 20 h shows an accumulation of 4N DNA content cells, corresponding to G2&M phase cells [39].

We also detect significant CCS signatures in experiments that have no apparent link to cell cycle arrest, direct or indirect. In a study comparing untransformed breast epithelial cells with breast cancer cell lines, three untransformed breast epithelial lines, MCF10A, HMT-3522 and HMEC1, showed significant correlation with one or more CCS. Cell lines were compared using a super-SILAC approach against MCF7, which is an ER+ breast cancer line. Both HMT-3522 and HMEC1 show strong correlation with early G1 states (CCS1). By contrast, MCF10A were correlated with S-phase (CCS4). Interestingly, ER+ MDA-MB-453 cultures also showed correlation with CCS1. These data suggest that the cell cycle distributions of these cell cultures are shifted compared to MCF7. In a separate study, 16 out of 62 lymphoblastoid cell lines (LCLs) analysed by proteomics to identify quantitative trait loci (QTL) were significantly CCS correlated (Fig. 6E). Interestingly, they were correlated in different states: 12 correlated with CCS1 and/or CCS2 (G1 phase) and the remaining four correlated with CCS5 (G2/early mitosis). These data suggest there is significant heterogeneity in cell cycle distribution, impacting at least 25% of the LCLs compared. How much of the heterogeneity in cell cycle state correlation observed has a genetic basis or is due to technical variation in cell culture handling will be important to assess.

## Discussion

A major challenge with the comprehensive analysis of proteomes from low cell number samples is sample preparation. An on-column load of 200 ng peptide, the equivalent to the protein content of approximately 2,000 TK6 cells, is sufficient material to obtain proteome coverage of >4,000 proteins with current instrumentation. Removal of detergents used to produce soluble cell extracts by use of membrane filters [40], organic precipitation (with or without the aid of magnetic beads) [41][42] or SDS-PAGE gel extraction are protocols involving several steps and repeated exposure to new plastic surfaces that introduce opportunities for non-specific peptide and protein adsorption. Here, we have presented a minimalistic approach for preparing cells for proteomics called the ‘in-cell digest’. Cells are fixed with formaldehyde and methanol to effectively trap them in biochemical states, then directly digested with trypsin and desalted prior to LC-MS/MS analysis.

We show that the in-cell digest enables reproducible and quantitative analysis of proteomes from 2,000 TK6 and MCF10A cells using AMPL analysis. The AMPL approach overcomes the low duty cycle of the Orbitrap Elite to enable proteome analysis with a sensitivity comparable with current instruments. Newer instrumentation with higher duty cycles, including the TIMS-TOF Pro and Exploris 480, is expected to enable conventional DDA and DIA analyses of proteomes at a similar depth with 2,000 TK6 cells, or alternatively, improve proteome depth further using MS1-based matching methods.

The in-cell digest is compatible with other approaches of low cell number sample preparation for MS-based proteomics. In-cell digested samples can be efficiently labelled by isobaric tags, e.g. TMT and iTRAQ, and therefore compatible with use of carrier channels to boost the signal of rare or single cell channels (e.g. iBASIL [43]). The protocol requires no specialized humidified sample handling chambers or direct loading onto premade analytical nanoLC columns, such as those described in the nanoPOTS workflow [11]. While the proteome coverages obtained by nanoPOTS is higher than reported here, it is possible that a new workflow combining aspects of the in-cell digest and nanoPOTS could improve both generalizability and performance compared to either method.

Each sample preparation method will have its unique advantages and potential biases, which we evaluated by quantitatively comparing the in-cell digest with a more conventional in-solution digest. This analysis revealed an overrepresentation of membrane proteins amongst those proteins with higher abundance measured in the in-cell digest samples. These proteins include mitochondrial membrane proteins (e.g. TOMM7) and proteins that are known to be localized to the cell surface (ADAM15). Membrane proteins have been shown to irreversibly aggregate in soluble extracts when heat-treated and precipitated. Delipidation by methanol, which is used to increase cell permeability, could also play an important role in increasing digestion efficiency of membrane proteins by trypsin. We suggest that the higher abundances measured for membrane proteins is unlikely to be an artefact of the in-cell digest; in contrast, the measurements are likely to more accurately reflect the abundances of these proteins in cells.

Feature matching FDR is controlled in our approach by implementing stringent cutoffs for retention time difference, m/z difference and match m/z error. Using a chemically modified ‘decoy’ proteome, we demonstrate that these cutoffs reduce the false positive rate with minimal impact on true positives. Elution time filtering provided greater discrimination between true and false positives than mass accuracy, suggesting that further improvements in chromatographic precision will benefit FDR control. We detect a higher estimated FDR compared to previous published models using mixed species [44]. However, our analysis differs in two significant aspects: 1) unlike matching between individual ‘single shot’ analyses, our experimental approach assesses match FDR from a fractionated library to a single shot analysis, and 2) unlike a mixed species proteome, our decoy proteome lacks true positives that could prevent assignment to false positive features. The latter means our reported FDR is likely an overestimate, but does provide a metric for assessing the relative FDR when filtering on feature match parameters. Additionally, models based on mixed-species suggest that matching FDR increases at low sample loads. It will be important in future to assess this with AMPL. In this study, comparable on-column loads between FDR estimation and cell cycle analysis and therefore, we are confident in the performance of false positive removal in the cell cycle dataset.

We identify novel proteins whose cell cycle function has not been previously characterized. FAM111B is a pseudoperiodic protein in cluster 1 (Fisher’s p_1_ < 0.001, p_2_ = 0.06), showing peak levels in S-phase populations, followed by a decrease in G2 populations. FAM111B is poorly characterized despite its expression being associated with poor prognosis in pancreatic and liver cancers (Human Protein Atlas [45]) and mutation causative for a rare inherited genetic syndrome (hereditary fibrosing poikiloderma with tendon contracture, myopathy, and pulmonary fibrosis). FAM111A, the only other member of the FAM111 gene family, localizes to newly synthesized chromatin during S-phase, interacts with PCNA via its PCNA-interacting protein (PIP) box and its depletion reduces base incorporation during DNA replication [46]. FAM111B also contains a PIP box (residues 607 – 616). Data from HeLa S3 cells also suggest that FAM111B is a cell cycle regulated protein with peak levels in S-phase [47]. FAM111B contains D-box and KEN-box motifs that are recognized by the APC/C E3 ligase to ubiquitinate targets for proteasomal degradation. Due to the similarity with FAM111A in sequence, predicted interactions with PCNA and peak protein abundance in S-phase, we propose that FAM111B also is likely to play a key role in DNA replication.

We present an unbiased pseudotemporal analysis of protein abundance changes across 8 biochemically resolved mitotic states with a resolution extremely challenging to obtain with high precision using arrest and release methodologies. The frequency of PsPs identified (1.7%, 119 / 6,899) compares well with a recent antibody-based screen for cell cycle regulated proteins (2.6%, 331 / 12,390) [48]. Included in 331 hits are proteins that vary in subcellular localization but not abundance across the cell cycle, consistent with other datasets using biochemical fractionation [49]. PsPs identified in this study will be limited to proteins that change in abundance. However, these PsPs are critical for robust cell state classification of proteomes obtained by MS, most of which do not involve subcellular fractionation.

A high proportion of proteins in clusters 4 and 5 (24/69, 35%) are experimentally validated APC/C substrates, which represents a 70-fold overrepresentation in these two clusters compared to non-pseudoperiodic proteins (0.5%). The high mitotic phase resolution and purity obtained in this study enabled characterisation of protein abundance variation of APC/C substrates in mitosis. We identify two waves of mitotic degradation, one coinciding with the destruction of cyclins A and B (cluster 4) and the second at mitotic exit (cluster 5). The unbiased clustering failed to separate cyclin A and cyclin B, which are degraded in prometaphase and at the metaphase-to-anaphase transition, respectively. This can be explained by the relatively few proteins detected that correlate with cyclin A and is consistent with the idea that prometaphase degradation by the APC/C is highly selective. 44 proteins in clusters 4 and 5 have not been previously experimentally validated as APC/C substrates [29] and are candidates for future follow-up analysis as novel, uncharacterized substrates. These include proteins (e.g. PRC1, KIF23, KIF20A) that were not identified as APC/C-Cdh1 and APC/C-Cdc20 substrates by bioinformatic analysis of co-regulation [50] and by chemical biology approaches [51,52].

High resolution classification of cell cycle state is an important prerequisite to obtaining meaningful biological insights into single cell ‘omics’ data. However, datasets on the cell cycle regulated transcriptome and proteome generally provide low time resolution, particularly in mitosis. Mitotic time resolution will be crucial for interpreting single cell proteomes. Whereas transcriptional and translational activity are dampened during mitosis, there are major changes in protein phosphorylation and protein abundance, which will contribute towards single cell proteome variation.

Here we have identified a robust cell cycle signature composed of the abundances from 119 PsPs that can be used to classify the cell cycle state of a cell population by virtue of its cellular proteome. We apply this signature to assign cell cycle states to hundreds of published proteomic datasets that range in cell type and experimental condition. We have not tested if this signature can be used to assign proteomes from species other than human. We note that many of these proteins are well conserved, with several conserved to yeast (e.g. cyclin, REC8, Aurora kinase, Polo kinase). We anticipate that this high-resolution cell cycle signature here will be important to understand the biological implications of emerging single cell proteomics datasets [9,10], particularly in systems where cell cycle phase differences are an underlying source of variation, as is frequently the case.

Formaldehyde fixation is used frequently as a precursor to intracellular immunostaining for cellular analysis and for inactivating cells that potentially harbor infectious agents, e.g. viruses. We have shown that mild formaldehyde treatment is compatible with comprehensive and quantitative proteomics with low cell numbers. We anticipate that the in-cell digest will be broadly applicable to characterise the proteomes of formaldehyde fixed, virally-infected cells. Recently published data suggest that formaldehyde crosslinks can be directly detected from MS data [53]. We anticipate the in-cell digest would enhance the sensitivity of crosslink detection and lead to an increase in identified protein-protein interactions.

Abbreviations: PRIMMUS, AMPL

## Acknowledgements

This work was supported by a Sir Henry Dale Fellowship to TL (Wellcome Trust & Royal Society 206211/Z/17/Z), a Darwin Trust PhD Studentship to ASA, an EASTBIO PhD Studentship to DAL, an MRC Career Development Award to GK, the Wellcome Centre for Cell Biology (WCB) core facilities (Wellcome Trust 203149), and funding for instrumentation, including equipment grants to the WCB Proteomics Core (091020) and the flow facilities. We thank valuable feedback and discussions with colleagues in the WCB and the University of Edinburgh, including Fiona Rossi (Scottish Centre for Regenerative Medicine), Christos Spanos (WCB) and Shaun Webb (WCB).

## Data Availability

Raw MS data and processed MaxQuant output files are available on ProteomeXchange/PRIDE. These data can be accessed using the project accession number PXD020006. Username: Password: AUSLTsvu.

## Supplementary Tables

Supplementary Table 1. Error tolerant search results

Supplementary Table 2. Raw MaxQuant proteomic output for the 16 cell cycle populations

Supplementary Table 3. Quantitative and pseudoperiodic analysis of the 16 cell cycle proteomes.

Supplementary Table 4. Pseudoperiodic proteins (PsPs) identified in this dataset. Supplementary Table 5. Short linear motif (SLiM) enrichment analysis.

Supplementary Table 6. 8-state cell cycle signature.

Supplementary Table 7. Application of cell cycle signature to ProteomeHD.

## Supplementary Data

Supplementary Data 1. R scripts used for the evidence FDR filtering, pseudoperiodicity analysis and proteomeHD analysis

## Supplementary Figures

**Supplementary Figure 1.**
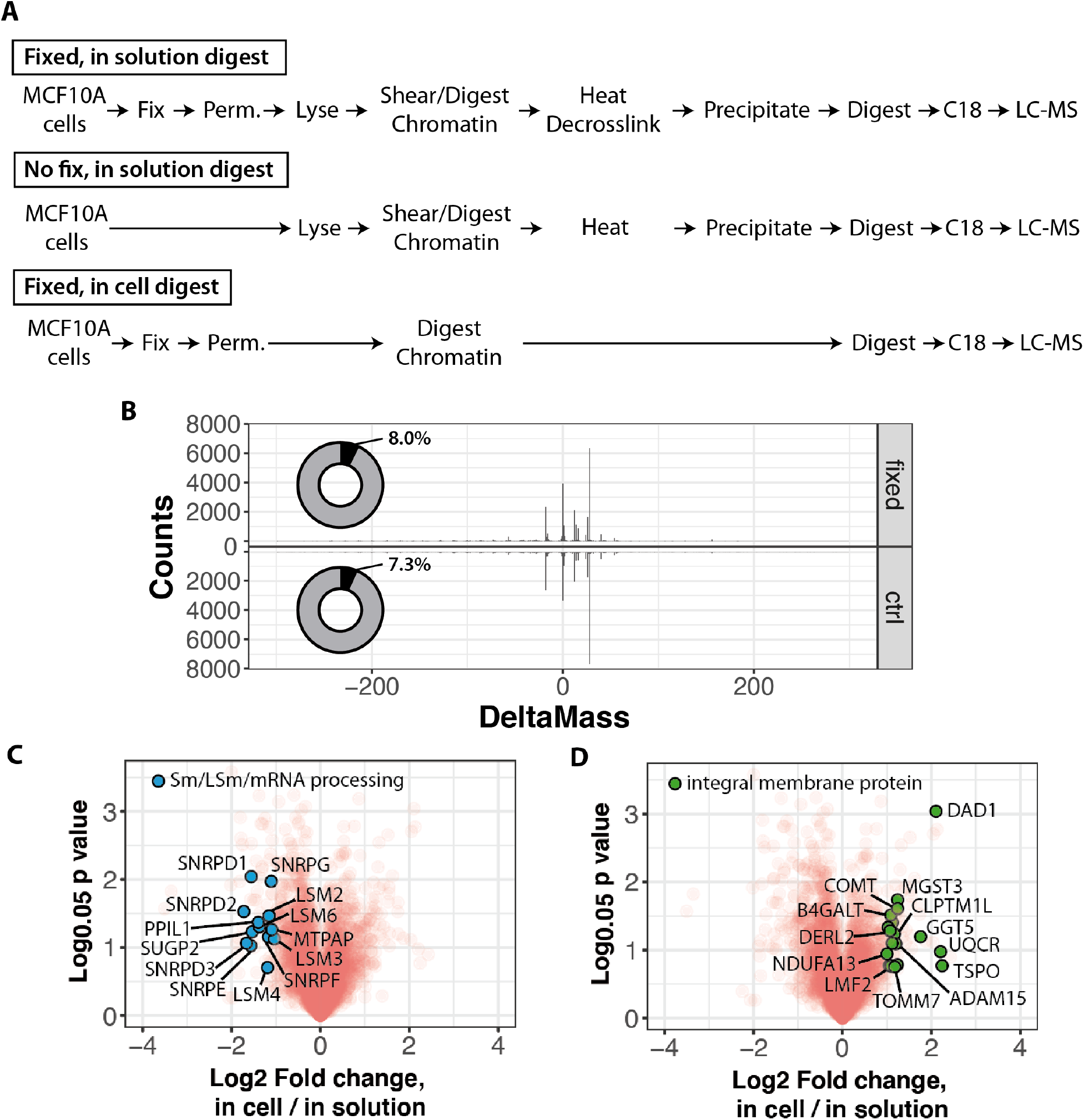
A) Scheme in Fig. 1A expanded to include details. B) Results from an error-tolerant dependent peptide search (MaxQuant) comparing control (cells without fixative) and fixed cells. C,D) Volcano plots comparing triplicate in cell and in solution digests.

**Supplementary Figure 2.**
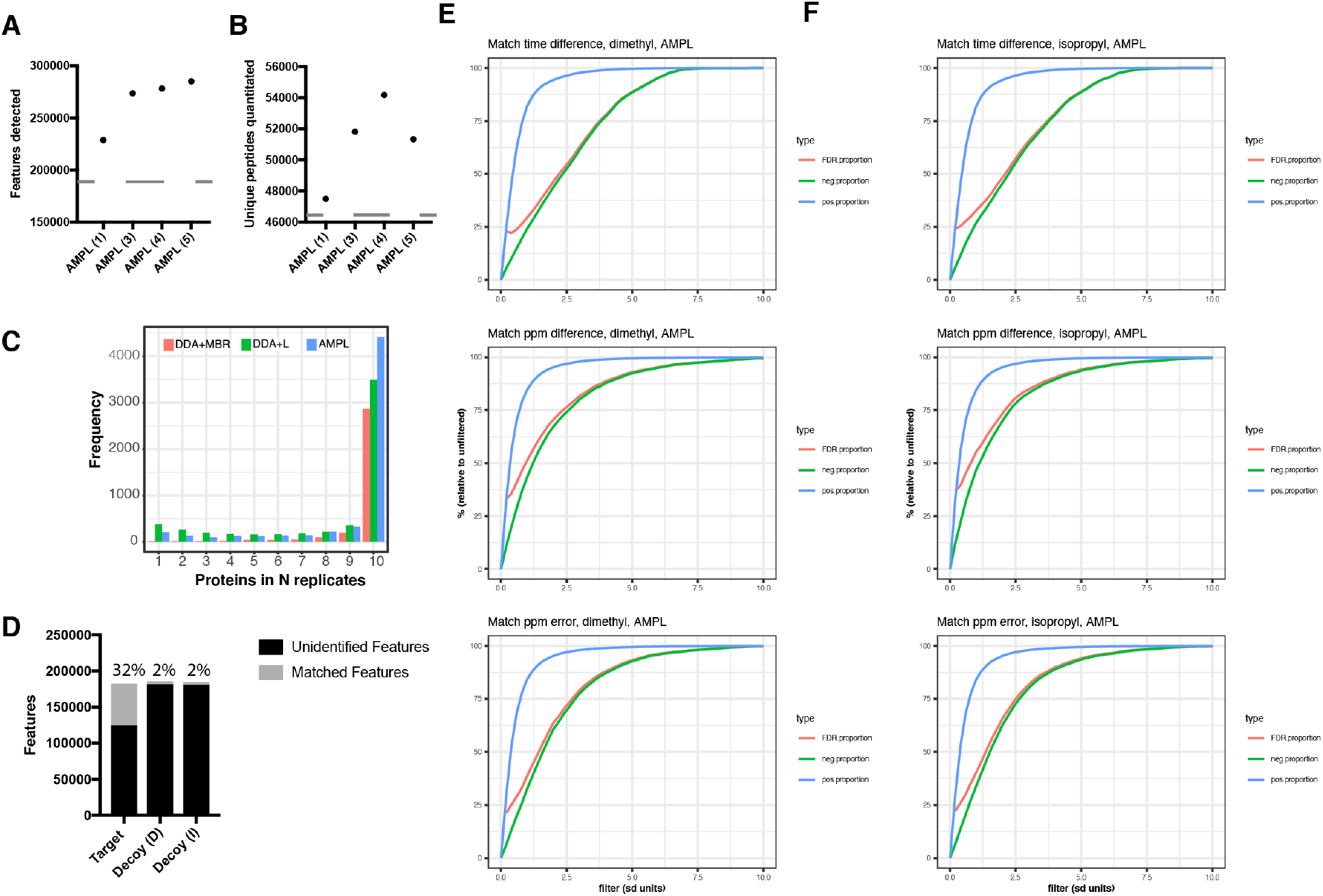
A) Identified features in target and decoy proteomes, including proportion of features matched to library. B, C) Optimising the number of MS1 spectra to average by measuring impact on the number of features and peptides detected. D) An analysis of data completeness comparing DDA, DDA with a library (DDA+L) and MPL. E, F) ROC plots for the dimethyl (E) and isopropyl (F) decoy proteomes filtering against match time difference, match ppm difference and match ppm error. Proportions are given in %s. FDR proportion is the estimated FDR at indicated filtering threshold over FDR with no filter.

**Supplementary Figure 3.**
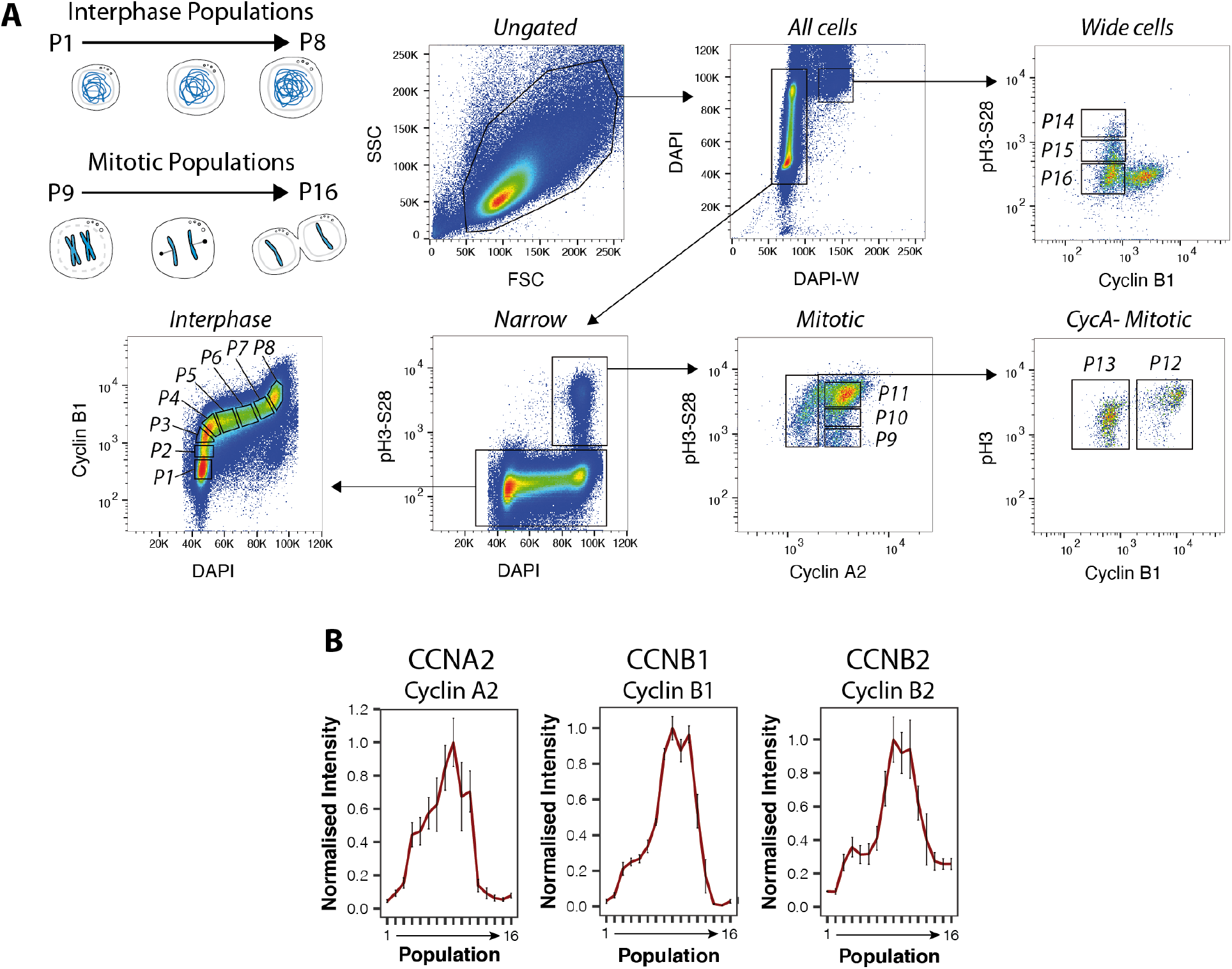
A) Scheme of FACS isolation and pseudocolour plots showing the gating strategy to isolate the 16 cell cycle populations. B) Mean normalised intensities across the 16 cell cycle populations as measured by MS for cyclin A2, cyclin B1 and cyclin B2.

## References

1. Ly T, Whigham A, Clarke R, Brenes-Murillo AJ, Estes B, Madhessian D, Lundberg E, Wadsworth P, Lamond AI (2017) Proteomic analysis of cell cycle progression in asynchronous cultures, including mitotic subphases, using PRIMMUS. eLife 6: e27574.

2. Ly T, Endo A, Lamond AI (2015) Proteomic analysis of the response to cell cycle arrests in human myeloid leukemia cells. eLife 4:.

3. Ly T, Whigham A, Clarke R, Brenes-Murillo AJ, Estes B, Wadsworth P, Lamond AI (2017) Proteomic analysis of cell cycle progression in asynchronous cultures, including mitotic subphases, using PRIMMUS. bioRxiv 125831.

4. Kelly RT (2020) Single-cell Proteomics: Progress and Prospects. Mol Cell Proteomics 19: 1739–1748.

5. Specht H, Emmott E, Petelski AA, Huffman RG, Perlman DH, Serra M, Kharchenko P, Koller A, Slavov N (2021) Single-cell proteomic and transcriptomic analysis of macrophage heterogeneity using SCoPE2. Genome Biol 22: 50–27.

6. Wiśniewski JR, Zougman A, Nagaraj N, Mann M (2009) Universal sample preparation method for proteome analysis. Nat Meth 6: 359–362.

7. Hughes CS, Foehr S, Garfield DA, Furlong EE, Steinmetz LM, Krijgsveld J (2014) Ultrasensitive proteome analysis using paramagnetic bead technology. Molecular Systems Biology 10: 757–757.

8. Zhang Z, Dubiak KM, Huber PW, Dovichi NJ (2020) Miniaturized Filter-Aided Sample Preparation (MICRO-FASP) Method for High Throughput, Ultrasensitive Proteomics Sample Preparation Reveals Proteome Asymmetry in Xenopus laevis Embryos. Anal Chem 92: 5554–5560.

9. Brunner A-D, Thielert M, Vasilopoulou CG, Ammar C, Coscia F, Mund A, Hoerning OB, Bache N, Apalategui A, Lubeck M, et al. (2021) Ultra-high sensitivity mass spectrometry quantifies single-cell proteome changes upon perturbation. bioRxiv 2020.12.22.423933.

10. Hartlmayr D, Ctortecka C, Seth A, Mendjan S, Tourniaire G, Mechtler K (2021) An automated workflow for label-free and multiplexed single cell proteomics sample preparation at unprecedented sensitivity. bioRxiv 2021.04.14.439828.

11. Zhu Y, Piehowski PD, Zhao R, Chen J, Shen Y, Moore RJ, Shukla AK, Petyuk VA, Campbell-Thompson M, Mathews CE, et al. (2018) Nanodroplet processing platform for deep and quantitative proteome profiling of 10–100 mammalian cells. Nat Comms 9: 882.

12. Metz B, Kersten GFA, Hoogerhout P, Brugghe HF, Timmermans HAM, de Jong A, Meiring H, Hove ten J, Hennink WE, Crommelin DJA, et al. (2004) Identification of Formaldehyde-induced Modifications in Proteins REACTIONS WITH MODEL PEPTIDES. Journal of Biological Chemistry 279: 6235–6243.

13. Toews J, Rogalski JC, Clark TJ, Kast J (2008) Mass spectrometric identification of formaldehyde-induced peptide modifications under in vivo protein cross-linking conditions. Anal Chim Acta 618: 168–183.

14. Skopek TR, Liber HL, Penman BW, Thilly WG (1978) Isolation of a human lymphoblastoid line heterozygous at the thymidine kinase locus: Possibility for a rapid human cell mutation assay. Biochemical and Biophysical Research Communications 84: 411–416.

15. Cox J, Mann M (2008) MaxQuant enables high peptide identification rates, individualized p.p.b.-range mass accuracies and proteome-wide protein quantification. Nat Biotechnol 26: 1367–1372.

16. Mohammed H, Taylor C, Brown GD, Papachristou EK, Carroll JS, D’Santos CS (2016) Rapid immunoprecipitation mass spectrometry of endogenous proteins (RIME) for analysis of chromatin complexes. Nat Protoc 11: 316–326.

17. Kong AT, Leprevost FV, Avtonomov DM, Mellacheruvu D, Nesvizhskii AI (2017) MSFragger: ultrafast and comprehensive peptide identification in mass spectrometry–based proteomics. Nat Meth 14: 513–520.

18. Pasa-Tolic L, Masselon C, Barry RC, Shen Y, Smith RD (2004) Proteomic analyses using an accurate mass and time tag strategy. BioTechniques 37: 621–639.

19. Meier F, Geyer PE, Winter SV, Cox J, Mann M (2018) BoxCar acquisition method enables single-shot proteomics at a depth of 10,000 proteins in 100 minutes. Nat Meth 15: 440–448.

20. Pines J, Clute P (1999) Temporal and spatial control of cyclin B1 destruction in metaphase. Nat Cell Biol 1: 82–87.

21. Elzen den N, Pines J (2001) Cyclin a Is Destroyed in Prometaphase and Can Delay Chromosome Alignment and Anaphase. The Journal of Cell Biology 153: 121–136.

22. Koo SJ, Fernández-Montalván AE, Badock V, Ott CJ, Holton SJ, Ahsen von O, Toedling J, Vittori S, Bradner JE, Gorjánácz M (2016) ATAD2 is an epigenetic reader of newly synthesized histone marks during DNA replication. Oncotarget 7: 70323–70335.

23. Kumar M, Gouw M, Michael S, Sámano-Sánchez H, Pancsa R, Glavina J, Diakogianni A, Valverde JA, Bukirova D, Čalyševa J, et al. (2019) ELM—the eukaryotic linear motif resource in 2020. Nucleic Acids Res 48: D296–D306.

24. Gonçalves V, Jordan P (2015) Posttranscriptional Regulation of Splicing Factor SRSF1 and Its Role in Cancer Cell Biology. Biomed Res Int 2015: 287048.

25. Breig O, Baklouti F (2013) Proteasome-mediated proteolysis of SRSF5 splicing factor intriguingly co-occurs with SRSF5 mRNA upregulation during late erythroid differentiation. PLoS ONE 8: e59137.

26. Maréchal A, Zou L (2013) DNA damage sensing by the ATM and ATR kinases. Cold Spring Harb Perspect Biol 5: a012716.

27. Kernan J, Bonacci T, Emanuele MJ (2018) Who guards the guardian? Mechanisms that restrain APC/C during the cell cycle. Biochimica et Biophysica Acta (BBA) - Molecular Cell Research 1865: 1924–1933.

28. Chang L, Barford D (2014) Insights into the anaphase-promoting complex: a molecular machine that regulates mitosis. Current Opinion in Structural Biology 29: 1–9.

29. Davey NE, Morgan DO (2016) Building a Regulatory Network with Short Linear Sequence Motifs: Lessons from the Degrons of the Anaphase-Promoting Complex. Molecular Cell 64: 12–23.

30. Di Fiore B, Davey NE, Hagting A, Izawa D, Mansfeld J, Gibson TJ, Pines J (2015) The ABBA Motif Binds APC/C Activators and Is Shared by APC/C Substrates and Regulators. Developmental Cell 32: 358–372.

31. Lara-Gonzalez P, Scott MIF, Diez M, Sen O, Taylor SS (2011) BubR1 blocks substrate recruitment to the APC/C in a KEN-box-dependent manner. Journal of Cell Science 124: 4332–4345.

32. Hellmuth S, Gómez-H L, Pendás AM, Stemmann O (2020) Securin-independent regulation of separase by checkpoint-induced shugoshin-MAD2. Nature 580: 536–541.

33. Stegle O, Teichmann SA, Marioni JC (2015) Computational and analytical challenges in single-cell transcriptomics. Nat Rev Genet 16: 133–145.

34. Scialdone A, Natarajan KN, Saraiva LR, Proserpio V, Teichmann SA, Stegle O, Marioni JC, Buettner F (2015) Computational assignment of cell-cycle stage from single-cell transcriptome data. Human Pluripotent Stem Cells 85: 54–61.

35. Liu Z, Lou H, Xie K, Wang H, Chen N, Aparicio OM, Zhang MQ, Jiang R, Chen T (2017) Reconstructing cell cycle pseudo time-series via single-cell transcriptome data. Nat Comms 8: 22.

36. Cuomo ASE, Seaton DD, McCarthy DJ, Martinez I, Bonder MJ, Garcia-Bernardo J, Amatya S, Madrigal P, Isaacson A, Buettner F, et al. (2020) Single-cell RNA-sequencing of differentiating iPS cells reveals dynamic genetic effects on gene expression. Nat Comms 11: 810.

37. Ly T, Ahmad Y, Shlien A, Soroka D, Mills A, Emanuele MJ, Stratton MR, Lamond AI (2014) A proteomic chronology of gene expression through the cell cycle in human myeloid leukemia cells. eLife 3: e01630.

38. Kustatscher G, Grabowski P, Schrader TA, Passmore JB, Schrader M, Rappsilber J (2019) Co-regulation map of the human proteome enables identification of protein functions. Nat Biotechnol 37: 1361–1371.

39. Fierro-Monti I, Echeverria P, Racle J, Hernandez C, Picard D, Quadroni M (2013) Dynamic impacts of the inhibition of the molecular chaperone Hsp90 on the T-cell proteome have implications for anti-cancer therapy. PLoS ONE 8: e80425.

40. Wiśniewski, J. R., Zougman, A., Nagaraj, N., & Mann, M. (2009). Universal sample preparation method for proteome analysis. Nature Methods, 6(5), 359–362. http://doi.org/10.1038/nmeth.1322

41. Hughes, C. S., Foehr, S., Garfield, D. A., Furlong, E. E., Steinmetz, L. M., & Krijgsveld, J. (2014). Ultrasensitive proteome analysis using paramagnetic bead technology. Molecular Systems Biology, 10(10), 757–757. http://doi.org/10.15252/msb.20145625

42. Batth, T. S., Tollenaere, M. X., Rüther, P., Gonzalez-Franquesa, A., Prabhakar, B. S., Bekker-Jensen, S., et al. (2019). Protein Aggregation Capture on Microparticles Enables Multipurpose Proteomics Sample Preparation. Molecular & Cellular Proteomics: MCP, 18(5), 1027–1035. http://doi.org/10.1074/mcp.TIR118.001270

43. Yu, F., Haynes, S. E., & Nesvizhskii, A. I. (2021). IonQuant Enables Accurate and Sensitive Label-Free Quantification With FDR-Controlled Match-Between-Runs. Molecular & Cellular Proteomics: MCP, 20, 100077. http://doi.org/10.1016/j.mcpro.2021.100077

44. Tsai, C.-F., Zhao, R., Williams, S. M., Moore, R. J., Schultz, K., Chrisler, W. B., et al. (2020). An Improved Boosting to Amplify Signal with Isobaric Labeling (iBASIL) Strategy for Precise Quantitative Single-cell Proteomics. Molecular & Cellular Proteomics: MCP, 19(5), 828.

45. Uhlen M, Zhang C, Lee S, Sjöstedt E, Fagerberg L, Bidkhori G, Benfeitas R, Arif M, Liu Z, Edfors F, et al. (2017) A pathology atlas of the human cancer transcriptome. Science 357: eaan2507.

46. Alabert C, Bukowski-Wills J-C, Lee S-B, Kustatscher G, Nakamura K, de Lima Alves F, Menard P, Mejlvang J, Rappsilber J, Groth A (2014) Nascent chromatin capture proteomics determines chromatin dynamics during DNA replication and identifies unknown fork components. Nat Cell Biol 16: 281–291.

47. Aviner R, Shenoy A, Elroy-Stein O, Geiger T (2015) Uncovering Hidden Layers of Cell Cycle Regulation through Integrative Multi-omic Analysis. PLoS Genet 11: e1005554.

48. Mahdessian D, Cesnik AJ, Gnann C, Danielsson F, Stenström L, Arif M, Zhang C, Le T, Johansson F, Shutten R, et al. (2021) Spatiotemporal dissection of the cell cycle with single-cell proteogenomics. Nature 590: 649–654.

49. Herr P, Boström J, Rullman E, Rudd SG, Vesterlund M, Lehtiö J, Helleday T, Maddalo G, Altun M (2020) Cell cycle profiling reveals protein oscillation, phosphorylation, and localization dynamics. Mol Cell Proteomics mcp.RA120.001938.

50. Franks JL, Martinez-Chacin RC, Wang X, Tiedemann RL, Bonacci T, Choudhury R, Bolhuis DL, Enrico TP, Mouery RD, Damrauer JS, et al. (2020) In silico APC/C substrate discovery reveals cell cycle-dependent degradation of UHRF1 and other chromatin regulators. PLoS Biol 18: e3000975.

51. Bakos G, Yu L, Gak IA, Roumeliotis TI, Liakopoulos D, Choudhary JS, Mansfeld J (2018) An E2-ubiquitin thioester-driven approach to identify substrates modified with ubiquitin and ubiquitin-like molecules. Nat Comms 9: 4776–15.

52. Manohar S, Yu Q, Gygi SP, King RW (2020) The Insulin Receptor Adaptor IRS2 is an APC/C Substrate That Promotes Cell Cycle Protein Expression and a Robust Spindle Assembly Checkpoint. Mol Cell Proteomics 19: 1450–1467.

53. Tayri-Wilk T, Slavin M, Zamel J, Blass A, Cohen S, Motzik A, Sun X, Shalev DE, Ram O, Kalisman N (2020) Mass spectrometry reveals the chemistry of formaldehyde cross-linking in structured proteins. Nat Comms 11: 3128.

